# Animal retrozymes are non-autonomous sequences of *Penelope*-like elements

**DOI:** 10.64898/2026.07.22.740005

**Authors:** Zhen Li, Isabelle Clavereau, Nicolas Pollet

## Abstract

Hammerhead ribozymes (HHRs) are small catalytic RNAs found across diverse life forms. In animal genomes, they can be encoded by genes organised in dispersed copies or in tandem genomic arrangements. These tandemly organised forms, known as Non-LTR retrozymes, were recently identified as a distinct group of non-autonomous retrotransposons, likely mobilised via a rolling-circle transposition mechanism and potentially involved in host transcriptome regulation. However, their evolutionary origins remain poorly understood. Here, we investigate the presence, genomic distribution, and possible origins of Non-LTR retrozymes across a broad range of vertebrate species. We find that these elements display a patchy phylogenetic distribution, notably absent from the Aves and Mammalia lineages. In species where they are present, retrozyme copy number, consensus length, and monomer proportion vary widely across species and retrozyme families, suggesting diverse amplification dynamics. Genomic mapping reveals a significant enrichment of Non-LTR retrozymes in intergenic regions and their exclusion from introns and exons, indicating selective pressure against genic insertion. Strikingly, the phylogenetic distribution of Non-LTR retrozymes coincides with that of Penelope-like elements (PLEs). Phylogenetic analysis further shows that the pLTR region of PLEs is closely related to Non-LTR retrozymes, supporting the hypothesis that Non-LTR retrozymes are non-autonomous derivatives of PLEs. Together, our findings shed new light on the evolutionary origin and genomic behaviour of Non-LTR retrozymes and underscore their potential regulatory roles in vertebrate genomes.

## Introduction

Eukaryotic genomes contain a vast array of repetitive elements, which underlie the C-value paradox: the tree of life and organisms’ genomic DNA content are unrelated (1). Although it was once considered ‘junk DNA’, increasing evidence suggests that many repetitive elements play significant roles in gene regulation and genome evolution (2–5). Among repetitive elements, transposable elements (TEs) are the most prominent group, as they have a strong ability to shape genomic structure and expression profiles (4). TEs are mobile DNA that can transpose into the host genome. Based on their transposition intermediates, TEs can be RNA TEs (Class I), which transpose via an RNA intermediate, or DNA TEs (Class II), which transpose via a DNA intermediate. Each class is further divided into superfamilies based on their detailed transposition mechanisms, but numerous repetitive sequences belong to the genomic dark matter and remain unclassified (6, 7).

The insertion and excision of TEs can disrupt cis-regulation in host genomes and alter gene expression by modifying typical gene structures(4, 5, 8). In addition to their prominent roles in shaping genome architecture, some non-LTR retrotransposable elements, such as long Interspersed Nuclear Elements (LINEs) and Penelope-like Elements (PLEs), can influence host transcriptomes through a ribozyme self-cleaving activity (9–12). Self-cleaving ribozymes are essential components of the RNA world and play important roles in biological processes such as gene regulation and immune response (13–15). Still, their contribution to the landscape of repetitive DNA remains relatively underexplored compared with other components of TE biology. In addition to these specific TEs, other repetitive elements, such as the recently discovered retrozymes, encode ribozymes (16, 17) (Figure 1). Retrozymes contain the hammerhead ribozyme (HHR), which is ubiquitous across the tree of life, including viroids, eukaryotes, and prokaryotes (18–24). HHRs can adopt one of three folding structures: type I, II, or III (25–27). Based on the HHR fold that retrozymes harbour and their sequence organisation, retrozymes have been classified into LTR and non-LTR retrozymes (16, 17). The LTR retrozyme is organised like LTR transposons, with long terminal repeats at both ends, and harbours the type III HHR at its LTRs (16) (Figure 1). In contrast, non-LTR retrozymes are organised into tandem repeats and harbour the type I HHR (17) (Figure 1). Additionally, LTR retrozymes are primarily found in plants, whereas Non-LTR retrozymes are mostly found in metazoans (17).

**Figure 1.**
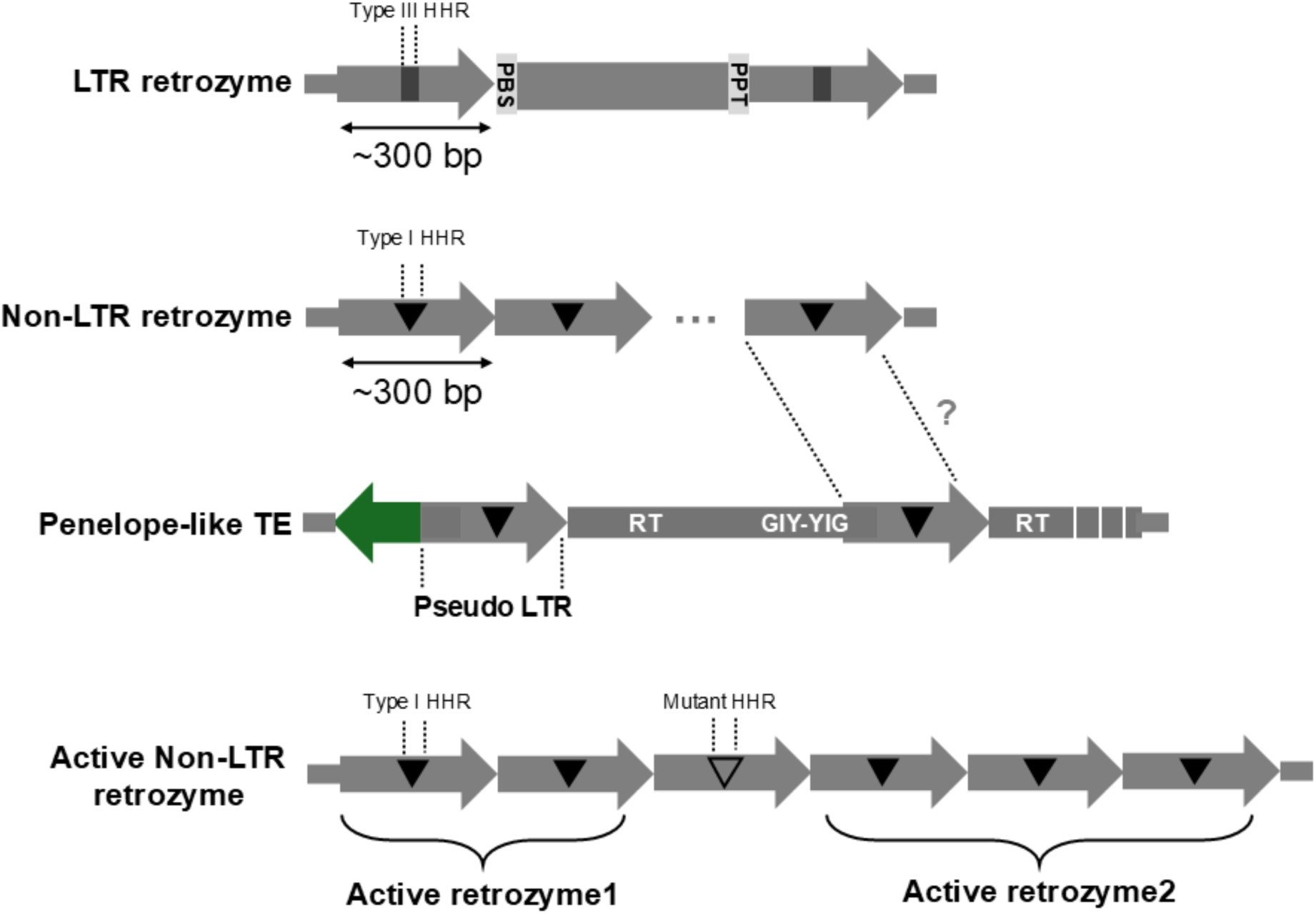
HHRs in retrozymes and PLEs. Retrozymes can be classified into two major types: LTR retrozymes and Non-LTR retrozymes. The LTR retrozyme contains the type III HHR in its LTR. The Non-LTR retrozyme contains a type I HHR in each tandem repeat unit. The type I HHR can also be detected within the pseudo-LTR of PLE retrotransposons. In this study, we defined an active Non-LTR retrozyme as an element containing the type I HHR in all its tandem repeats.

In previous studies, the transcription, self-cleavage and formation of circular RNA molecules from active retrozymes have been reported (16, 17, 25, 28). Similar to certain viroids and viral satellites, such circular RNAs are believed to replicate via a rolling-circle mechanism (16, 17, 25). Subsequently, they may be reverse-transcribed and inserted into new chromosomal loci (16, 17, 25). Their possible ability to transpose into the host genome makes them mobile genetic elements. Actually, both types of retrozymes don’t encode any proteins and are thought to be non-autonomous retrotransposons (16, 17, 25). The LTR retrozymes share a small conserved region of the LTRs of Gypsy retrotransposons, i.e., primer binding sites (PBS) and polypurine tract (PPT) (16). In addition, LTR retrozymes are flanked by 4-bp target-site duplications (TSDs), as are LTR retrotransposons (16). These features suggest that the LTR retrozymes might be non-autonomous LTR retroelements (16). The Non-LTR retrozymes harbour the type I HHR, which are also found in the pseudo LTRs of PLE retrotransposons, suggesting the Non-LTR retrozymes might originate from PLEs (17). Yet, the relationships between retrozymes and retrotransposable elements have not been investigated at the phylogenetic level.

Here, we focused on Non-LTR retrozymes and investigated their distribution across divergent metazoans. We comprehensively examined the phylogenetic relationship between the Non-LTR retrozymes and PLEs. Our work provides strong evidence that the Non-LTR retrozymes in metazoan host genomes originate from PLEs.

## Materials and methods

### Detection of retrozyme sequences from genome sequences

We downloaded the genomes of 35 animal species from NCBI, including 31 vertebrates and four invertebrates (Supplementary Table 1). We developed a Python-based pipeline to detect non-LTR retrozyme sequences from sampled genomes. In the first step, we detected tandem repeats using the trf program (v4.09) with the parameters “2 7 7 80 10 50 1000 -l 6 -h -ngs” (29). Next, we applied rnabob (v2.2.1) to search for HHR motifs within each tandem repeat, using both canonical and minimal type I HHR models (30). For each HHR sequence, we used RNAfold (v2.6.4) with default parameters to estimate its minimum free energy (MFE) (31). HHR sequences with an MFE lower than minus five kcal/mol were considered active. We considered tandem repeats containing at least two consecutive units with complete HHR motifs as active retrozymes (Figure 1). We based our criterion on the known biological requirement that active retrozymes require two HHR motifs to initiate their transposition cycle (15).

### Construction of consensus sequences for Non-LTR retrozymes

For each species, we classified the non-LTR retrozymes into families based on sequence homology (90% identity) using VSEARCH (v2.22.1) with the parameters “-- id 0.9 --strand both --clusterout_id --consout” (32). This clustering process also generated a consensus sequence for each retrozyme family.

### Distribution of Non-LTR retrozyme sequences in genomes

For each non-LTR retrozyme family detected in a given genome, we used blastn (v2.15.0, parameter: “-evalue 1e-5”) to search for full-length homologous sequences, using the family consensus sequence as the query (33). We defined a full-length homologous sequence as one sharing more than 80% identity over at least 80% of the consensus sequence length. We categorised these full-copy retrozyme sequences into monomers, dimers, and multimers based on their sequence organisation. The monomer referred to the “full-length” sequence that appeared as “solo retrozymes”; Dimers referred to tandem repeats formed by two “full-length” sequences; Multimers referred to tandem repeats formed by more than two “full-length” sequences.

To investigate the distribution of Non-LTR retrozyme sequences in a targeted genome, we made 100 kbp sliding windows with a 50-kbp step size for each chromosome. We estimated the retrozyme sequence coverage per window.

We further analysed the distribution of retrozymes across genomic features, including intronic, exonic, and intergenic regions. For each retrozyme family, we used bedtools intersect (v2.30.0) to compare the coordinates of full-length retrozyme annotations with those of various genomic features (34). A retrozyme full-length region was assigned to a specific genomic feature if more than 50% of its sequence overlapped with that feature. We performed this analysis only for the two *Xenopus* species, *X. tropicalis* and *X. laevis,* as they are established model species with higher-quality genome annotations than those of other sampled species.

### Detection of PLE transposase sequences for sampled species

Previous studies of PLE phylogeny have shown that PLEs belong to one of eight major clades based on homology of reverse transcriptase (RT) sequences: Athena, Coprina, Penelope/Poseidon, Neptune, Nematis, Hydra, Naiad, and Chlamys (35). These clades can be further divided into two groups based on the presence or absence of the GIY-YIG endonuclease (EN): 1) EN minus group, which lacks the EN domain and includes the Athena and Coprina clades; 2) EN plus group, which contains the EN domain and includes the Penelope/Poseidon, Neptune, Nematis, Hydra, Naiad and Chlamys clades. In this study, we focus only on the EN plus PLEs because the EN minus PLEs were not found in vertebrates and are restricted to specific lineages (35).

We retrieved PLE transposase sequences from the Repbase database (v27.04) (36). We separated them into RT and GIY-YIG domains using HMMER’s hmmsearch (v3.3), with HMM models trained from the Conserved Domain Database (CDD): cd00304 for the RT domain and cd10442 for the GIY-YIG domain (37, 38). Then, we built HMM profiles for each PLE clade and each domain (RT and GIY-YIG domains) using HMMER’s hmmbuild (v3.3). For each genome, we predicted protein-coding sequences using getorf (EMBOSS:6.6.0.0) with the parameter “-minsize 200”(39). After, we searched for the RT and GIY-YIG EN domains using hmmsearch (-E 1e-5) with the trained HMM models. To identify PLE transposase sequences containing both RT and GIY-YIG EN domains, we used bedtools window with the parameters “-w1500 -sm - sw”. We defined a transposase region by the co-occurrence of both domains within 1,500 bp of each other. We did not impose a specific domain order, as previous studies have shown that PLE architectures are variable, with the GIY-YIG EN domain occurring either upstream or downstream of the RT domain.

### Phylogenetic analysis for the sequence of pLTR and non-LTR retrozymes

We retrieved pLTR regions from the Repbase collection for *X. tropicalis* and *X. laevis*. We then performed multiple sequence alignment of the pLTR sequences with the *Xenopus* Non-LTR retrozyme consensus sequence using MAFFT (v7.526) with default parameters (40). Then, we constructed the phylogenetic tree using phyml with the parameter “-a e”(41). Finally, we visualised the tree using ITOL (42).

### Experimental analysis of retrozyme activity

We designed primers to amplify nine predicted retrozyme elements by PCR from three Xenopus species (*X. borealis, X. tropicalis, X. laevis*). PCRs were run using 50 ng of genomic DNA from the appropriate species, 0.5 µM each of the forward and reverse primers, 2.5 mM each dNTP, 5 U of OneTaq HotStart polymerase (New England Biolabs), and the corresponding buffer, in a final reaction volume of 25 µL, with 35 amplification cycles. We checked PCR products by routine gel electrophoresis and further validated the amplicons by sequencing with the EXP-NBD103 and SQK- LSK109 kits on a Flongle flow cell (Oxford Nanopore Technologies). Each PCR product (20 µL) was purified using an equivalent volume of magnetic beads (NucleoMag, Macherey-Nagel), along with a positive control for the in vitro transcription reaction. In Vitro transcription was performed in a 25 µL final volume using 200 nM purified PCR product, 20 U of T7 RNA polymerase (Thermo), 6 mM MgCl2, 1 mM NTPs, 1X IVT reaction buffer (Roche), and 0.25 U of RNase Inhibitor. We took a 5 µL sample after 1, 5, 30, and 60 min of incubation at 37°C, then added 2 µL of 0.5 mM EDTA to stop the reaction. The products were analysed by capillary electrophoresis using the Experion RNA StdSens Analysis kit (Bio-Rad) with 1 µL of the reaction product, following the manufacturer’s instructions.

## Results

### Occurrences of retrozymes in vertebrate genomes

It has been shown that retrozymes in the amphibian *Ambystoma mexicanum* genome belong specifically to the Non-LTR retrozyme class, which forms tandem repeats with extremely high copy numbers((16). However, the knowledge of the distribution of these Non-LTR retrozymes in other vertebrates remained incomplete. Here, we investigated the occurrence of Non-LTR retrozymes in 31 sampled vertebrate genomes, including four Aves, five Reptilia, six Mammalia, seven Amphibia, one Sarcopterygii, seven Actinopterygii, and one Chondrichthyes (Figure 2). We also included three Insecta and one Anthozoa genomes as outgroups. These vertebrate genomes spanned a wide range of sizes, from the smallest (*Takifugu rubripes*, 0.38 Gbp) to the largest (*A. mexicanum*, 28.21 Gbp) (Figure 2).

**Figure 2.**
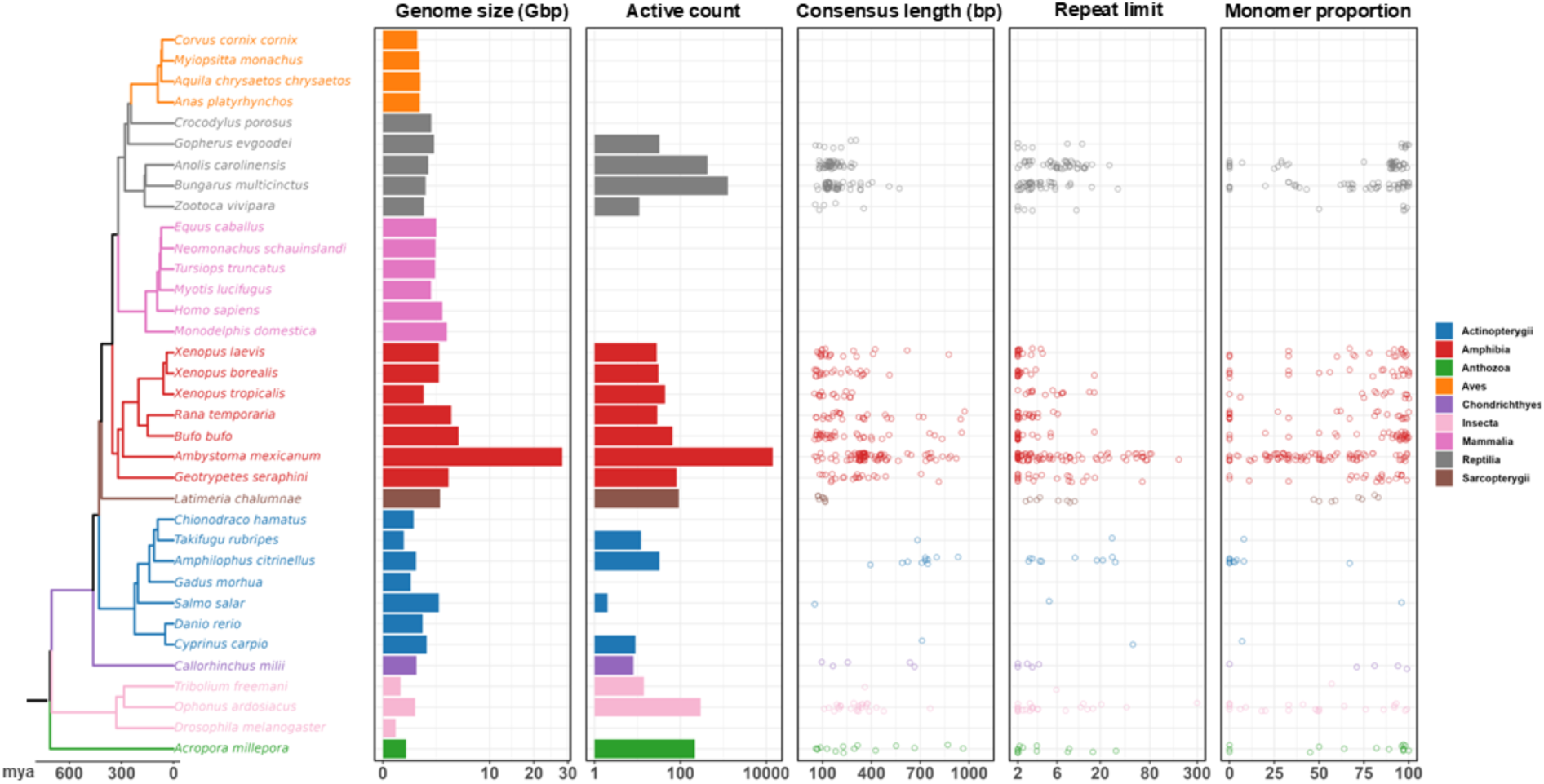
Occurrence of Non-LTR retrozymes in sampled vertebrates. The chronogram depicts the sampled vertebrates, with species from different taxonomic classes distinguished by colour. The figure is divided into five panels. 1) Genome size: Bar plot showing genome sizes in gigabase pairs (Gbp). 2) Active count: Bar plot displaying the number of active retrozymes in each genome. 3) Consensus length: Jitter plot showing the variation in the maximum consensus length across retrozyme families. 4) Repeat limit: Jitter plot indicating the maximum number of repeat units for each retrozyme family. 5) Monomer proportion: Jitter plot representing the variation in percentage of monomeric retrozymes relative to the total number of retrozyme repeat units across retrozyme families.

We applied a dedicated script to search for the Non-LTR retrozymes in each sampled species (Supplementary Table 2). For each species, we estimated the number of putatively active Non-LTR retrozymes (active count, Figure 2). We found Non-LTR retrozymes in 20 out of 31 vertebrate, 2 out of 3 insect, and 1 out of 1 Anthozoa genomes, with a maximum of 14,255 copies in the largest genome (*A. mexicanum*) and a minimum of 2 copies in the *Salmo salar* genome (Figure 2). Yet, we found 12 copies in the smallest genome (*T. rubripes*). We observed that the four Aves and six Mammalia classes were devoid of Non-LTR retrozymes. But these retrozymes were present in other sampled vertebrate classes, including four out of five Reptilia, all seven Amphibia, one Sarcopterygii, five out of seven Actinopterygii, and one Chondrichthyes species (Figure 2).

In genomes containing Non-LTR retrozymes, we classified retrozyme sequences into families. We found that the length of retrozyme consensus sequences varied across families from 51 to 972 bp, with averages of 316 bp in Amphibia, 650 bp in Actinopterygii, 176 bp in Reptilia, 98 bp in Sarcopterygii, 363 bp in Chondrichthyes, 342 bp in Anthozoa, and 313 bp in Insecta. For each retrozyme family, we recorded the maximum number of repeat units (Figure 2, Repeat limit). The result showed that the Non-LTR retrozyme could reach high repeat numbers, e.g. 300 repeat units in the *Ophonus ardosiacus* genome and 180 in the *A. mexicanum* genome. But the majority (75%) of retrozyme families had a maximum of seven repeat units. Furthermore, we investigated the sequence organisation of different retrozyme families by evaluating the proportion of their monomer derivatives, which might correspond to the offspring of active retrozymes that could not transpose. We found that the proportion of monomers varied among retrozyme families and species, ranging from 0 to 1.

We concluded that Non-LTR retrotransposons are pervasive in vertebrates, with only a few classes lacking them, such as Aves and Mammalia. Moreover, the amplification pattern of Non-LTR retrozymes varied greatly among different retrozyme families and species.

### The genomic distribution of Non-LTR retrozymes

Previous studies have shown that tandem repeats are enriched in telomeric and centromeric regions. We thus asked if the distribution pattern of retrozymes aligned with that of tandem repeats. We used the genome of the model species *X. tropicalis* as a test organism because it has better genome annotation than that of non-model organisms((43). For each Non-LTR retrozyme family, we searched the genome for homologous sequences using its consensus sequence as a query. These homologous sequences were found as monomers, dimers or multimers. We discovered that monomer sequences were evenly distributed across each chromosome, whereas dimer and multimer sequences were enriched in a few genomic regions (Figure 3A). In contrast, we observed that multimer sequences were enriched at the termini or centres of chromosomes, suggesting enrichment in telomeric and centromeric regions (Figure 3A).

**Figure 3.**
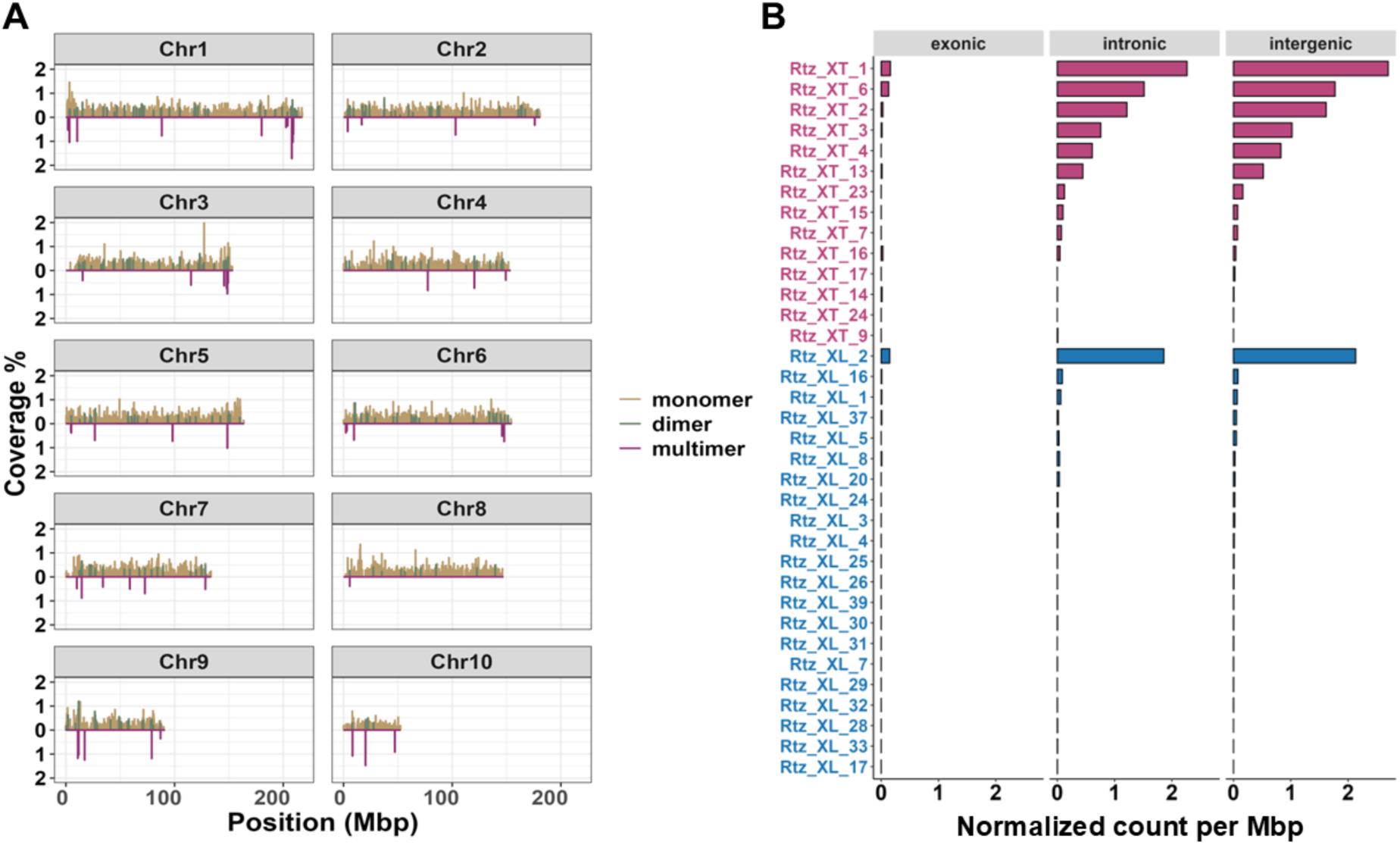
The distribution of Non-LTR retrozyme in genomes of *Xenopus* frogs. A: The distribution of retrozyme homologous sequences across chromosomes of *X. tropicalis*. These homologous regions were classified into three types: monomer, dimer (tandem repeats of two units), and multimer (tandem repeats of more than two units). B: Bar plot showing the normalised copy number of retrozyme homologous sequences in genomic features of *X. tropicalis* (in red) and *X. laevis* (in blue).

In vertebrates, TE insertions are typically missing in exons and do not alter standard transcriptional unit structure, although a few insertions within genes can contribute to the emergence of new exons (44). We analysed the distribution of non-LTR retrozymes across genomic features in the model species *X. tropicalis* and *X. laevis*, categorising the genome into three central regions: intergenic, exonic, and intronic (Supplementary Table 3). In *X. tropicalis*, we found that the majority (55.85%) of Non-LTR retrozymes overlapped the intergenic region, followed by 43.91% overlapping the intronic region and 0.24% overlapping the exonic region. We observed a similar pattern in the *X. laevis genome*, with 57.40%, 42.27%, and 0.32% overlap with the intergenic, intronic, and exonic regions, respectively. Since the three genomic features vary in size, we normalised the counts by each feature’s size. Still, we found that the intergenic region had the most abundant retrozyme copies, followed by the intronic region (Figure 3B). We discovered that retrozyme sequences tend to be enriched in the intergenic region and avoided in the intronic and exonic regions (Fisher’s exact test, p-value < 1.3e-18). In addition, we found that the retrozymes overlapping exons tended to be located at the last exon, more specifically, the 3’UTR region (Supplementary Table 4).

### PLE origin of the Non-LTR retrozymes

Previous studies have shown that type I HHRs are present in both Non-LTR retrozymes and terminal non-coding regions of PLEs, suggesting a close relationship in their biogenesis. We aimed to identify phylogenetic evidence for the relationship between the two elements.

We investigated the PLE distribution across all sampled species by searching for PLE transposase sequences using a homology-based approach. We observed PLEs in 24 of 35 species, belonging to five vertebrate and two invertebrate classes (Supplementary Table 5). We did not find any PLE sequences in Aves and Mammalia (Figure 4A). According to the PLE classification proposed in a previous study, we classified the detected PLEs into four major groups: Hydra, Naiad, Neptune, and Poseidon (35, 45). Of the four groups, Neptune and Poseidon were the most widespread, occurring in five vertebrate and two invertebrate classes. At the same time, Hydra and Naiad were only detected in the coral genome (*Acropora millepora*). We compared the phylogenetic distributions of PLEs and Non-LTR retrozymes and found that the two elements showed the same distribution in 32/35 species, indicating their co-occurrence (Figure 4A). The 22 species that contain Non-LTR Retrozymes also contain PLEs. Still, four species containing PLEs did not contain Non-LTR retrozymes, suggesting that the biogenesis of Non-LTR retrozymes might rely on the presence of autonomous PLEs (Figure 4A).

**Figure 4.**
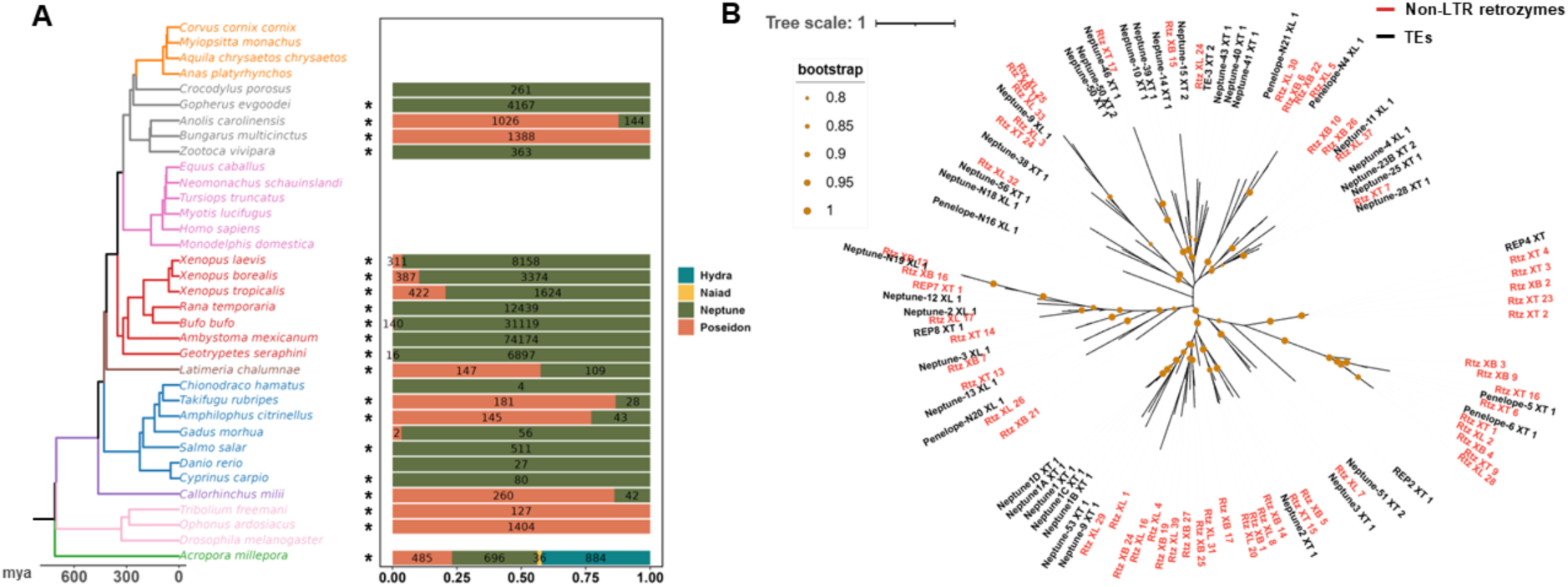
The phylogeny relationship between Non-LTR retrozymes and PLEs. A: The content of PLEs in sampled species. Hydra, Naiad, Neptune, and Poseidon are different PLE variants. The asterisk marks co-occurrences of PLEs and Non-LTR retrozymes. B: The phylogenetic tree of Non-LTR retrozymes and pLTR of PLEs in *Xenopus* frogs. Non-LTR retrozymes were labelled in red. XT: *X. tropicalis*; XL: *X. laevis*; XB: *X. borealis*.

We then asked whether the Non-LTR retrozyme showed any homology to known PLE sequences. We conducted this investigation of the *Xenopus* frog genome because *X. tropicalis* and *X. laevis* have well-annotated PLE sequences. We searched the retrozyme consensus sequences against the Repbase collection. Of the 57 *Xenopus* retrozyme families, 24 were homologous to 22 TEs, with 21 being annotated as PLEs and one being annotated as REP4_XT (Supplementary Table 6). Notably, the regions of similarity between retrozymes and PLEs were found at either the left or right termini of the PLEs, corresponding to their pLTR regions. This suggests that the pLTRs of PLEs share sequence similarity with Non-LTR retrozymes. To further explore this relationship, we constructed a phylogenetic tree using the pLTRs of PLEs and all Non-LTR retrozyme consensus sequences from *Xenopus* frogs. The result confirmed that the *Xenopus* Non-LTR retrozymes were phylogenetically closely related to the pLTR of PLEs. In fact, nearly every retrozyme family appeared as a sister group to one or more PLE sequences in the tree (Figure 4B), supporting a shared evolutionary origin.

In summary, we observed the co-occurrence of PLEs and Non-LTR retrozymes in our sampled species. In addition, we demonstrated in *Xenopus* genomes that the sequences of Non-LTR retrozymes were phylogenetically related to PLEs.

### Experimental validation of the activity of Non-LTR retrozymes in vitro

We set out to challenge our bioinformatic predictions and test experimentally the self-cleaving activity of representative dimer elements from *X. borealis*, *X. laevis,* and *X. tropicalis*. We designed nine primer pairs to amplify retrozyme dimer elements from different families in one of the three Xenopus species, and obtained the expected sequence products in each case. We then performed in vitro transcription in the presence of 6 mM MgCl2 to monitor cis-cleavage activity after capillary electrophoresis. We observed a very high cleavage activity of one REP4 dimer element from *X. tropicalis*. In contrast, we found no cleavage product from the eight other elements, including REP4 from *X. borealis* or *X. laevis*.

## Discussion

In this study, we investigated the occurrence, genomic distribution, and possible evolutionary origin of Non-LTR retrozymes across a broad range of vertebrate species. Our analyses revealed that these elements exhibit a patchy phylogenetic distribution, being absent from major clades such as Aves and Mammalia, but present in a diverse set of other vertebrates. This patchy distribution suggests either multiple independent losses or a limited number of ancestral insertions followed by clade-specific expansions and extinctions. In comparison with the known vertebrate TE landscape, the Non-LTR retrozymes distribution resembles that of PLEs but also of Helitron-like elements (HLEs), with the notable exception that even bat genomes seem devoid of Non-LTR retrozymes. Most families of TEs, including PLEs, were likely represented in the last common ancestor of vertebrates, assuming a model in which vertical inheritance was the dominant mode of transmission (46). In addition, horizontal transfer of TEs (HTTs) likely played a significant role in shaping the abundance and distribution of TEs observed in vertebrate lineages by repeatedly influencing TE activity and transposition rates and by altering host-mediated defences (47, 48). Among vertebrates, the HTT pattern was frequently observed in amphibians, squamates, and ray-finned fishes. Therefore, we hypothesise that HTT may be an important factor in explaining the distribution pattern of PLEs, which in turn led to the appearance of Non- LTR Retrozymes (48).

The evolutionary dynamics of PLEs and Non-LTR Retrozymes are likely to follow different routes once an active PLE lands in a given genome. Because PLEs are autonomous TEs, they can insert into different chromosomal loci, but most insertions suffer from 5’ truncations and therefore contain a single terminal repeat (45). Once a PLE is inserted at a given locus, its terminal sequence could give rise to an independent Non-LTR Retrozyme that will follow its own dynamics of local amplification to give rise to tandem repeats. This scenario resembles the one described for MITEs derived from class II TEs (51). Early on, the match between portions of PLEs and repetitive satellite DNAs was noted (45, 49). Yet, the case of microsatellite birth (typically less than 10 bp motif size and originating from replication slippage and recombination) should not be confounded with that of minisatellites (up to several hundred bp motif size), such as those made of Non-LTR retrozyme sequences. The acknowledged model of satellite birth includes the formation of a tandem repeat, followed by expansion by unequal crossover (50). Altogether, this provides further empirical evidence of the interplay between satellite and TE sequences, the two main components of repetitive DNA in eukaryotic genomes (52).

We observed the high variability in retrozyme copy number, consensus length, and monomer proportion across species and retrozyme families. This diversity implies lineage-specific differences in amplification dynamics, possibly influenced by transposition activity, host genome defence mechanisms, or ecological and evolutionary pressures. The enrichment of retrozymes in intergenic regions and their exclusion from the genic areas further supports the idea that insertion into functional genomic loci may be deleterious, leading to purifying selection against such events.

The overlap between the distributions of Non-LTR retrozymes and PLEs, coupled with the phylogenetic proximity between the pLTR regions of PLEs and Non-LTR retrozymes, suggests a potential evolutionary relationship. These observations support a model in which Non-LTR retrozymes are non-autonomous elements that hijack the enzymatic machinery of PLEs to mobilise them. This is reminiscent of other well-known non-autonomous partnerships in the TE world, such as SINEs and LINEs (53). The presence of hammerhead ribozyme motifs in Non-LTR retrozymes may further contribute to their mobility and regulatory potential. However, the precise mechanisms by which they function remain to be elucidated (17).

Recent studies have highlighted the role of TEs in shaping gene regulation, chromatin organisation, and genome evolution. The potential involvement of Non-LTR retrozymes in host transcriptome regulation, possibly through the production of circular RNAs or regulatory transcripts, warrants further investigation. Functional studies will be necessary to determine whether these elements have been co-opted into regulatory networks or exert influence via RNA interference, splicing modulation, or other mechanisms.

Despite the insights provided, our study has limitations. We mainly focused on vertebrates, and we did not investigate whether Non-LTR retrozymes from other organisms are phylogenetically close to PLEs. In addition, we could only provide limited empirical evidence of the life cycle of these elements.

In summary, our work sheds light on the distribution, diversity, and potential evolutionary origin of Non-LTR retrozymes in vertebrates. We propose that they are likely non-autonomous derivatives of PLEs. Future research should aim to characterise their mobilisation mechanisms, interaction with PLEs, and functional impacts on host transcriptomes.

## Acknowledgements

We thank the members of the pole genome for their constructive feedback on this project.

## Funding

China Scholarship Council–Université Paris Saclay PhD fellowship [202106760020 to Z.L.].

## Conflict of interest disclosure

The authors declare that they have no financial conflict of interest with the content of this article. Nicolas Pollet is one of the PCI Genomics recommenders. The authors declare that they have no other competing interests.

## Data, script and code availability

Data and tools to prepare all supplementary data are available at https://github.com/Zhenlisme/retrozyme

## Supplementary information

Supplementary materials are available in the zenodo open archive under the DOI 10.5281/zenodo.21493920.

## Notes

Present address: Zhen Li; Max Planck Institute of Biology Tübingen; Max-Planck-Ring 5; 72076 Tübingen; Germany.

## Declarations

### Ethics approval and consent to participate

#### Author’s contributions

All authors read and approved the final manuscript. ZL: Data curation; Formal Analysis; Investigation; Methodology; Validation; Visualization; Writing – original draft; Writing – review and editing. IC: Investigation; NP : Conceptualization ; Data curation; Funding Acquisition; Investigation; Methodology; Project Administration; Supervision; Validation; Visualization; Writing – review and editing.

## Notes

https://github.com/Zhenlisme/retrozyme

### Competing Interest Statement

The authors have declared no competing interest.

### Summary of Updates

The entire manuscript has been modified to add line numbering to ease the review process.

## References

1. Gregory, T.R. (2005) CHAPTER 1 - Genome Size Evolution in Animals. In Gregory, T.R. (ed), The Evolution of the Genome. Academic Press, Burlington, pp. 3–87.

2. Belyayev, A., Josefiová, J., Jandová, M., Mahelka, V., Krak, K. and Mandák, B. (2020) Transposons and satellite DNA: on the origin of the major satellite DNA family in the Chenopodium genome. Mobile DNA, 11, 20.

3. Garrido-Ramos, M.A. (2017) Satellite DNA: An Evolving Topic. Genes, 8, 230.

4. Cosby, R.L., Chang, N.-C. and Feschotte, C. (2019) Host–transposon interactions: conflict, cooperation, and cooption. Genes Dev., 33, 1098–1116.

5. Transposable elements promote the evolution of genome streamlining 10.1098/rstb.2020.0477.

6. Wells, J.N. and Feschotte, C. (2020) A Field Guide to Eukaryotic Transposable Elements. Annu Rev Genet, 54, 539–561.

7. Wicker, T., Sabot, F., Hua-Van, A., Bennetzen, J.L., Capy, P., Chalhoub, B., Flavell, A., Leroy, P., Morgante, M., Panaud, O., et al. (2007) A unified classification system for eukaryotic transposable elements. Nat Rev Genet, 8, 973–982.

8. Du, A.Y., Chobirko, J.D., Zhuo, X., Feschotte, C. and Wang, T. (2024) Regulatory transposable elements in the encyclopedia of DNA elements. Nat Commun, 15, 7594.

9. Cervera, A. and De la Peña, M. (2014) Eukaryotic Penelope-Like Retroelements Encode Hammerhead Ribozyme Motifs. Molecular Biology and Evolution, 31, 2941–2947.

10. Eickbush, D.G. and Eickbush, T.H. (2010) R2 Retrotransposons Encode a Self-Cleaving Ribozyme for Processing from an rRNA Cotranscript. Mol Cell Biol, 30, 3142–3150.

11. Kläge, D., Müller, E. and Hartig, J.S. (2024) A comparative survey of the influence of small self-cleaving ribozymes on gene expression in human cell culture. RNA Biol, 21, 1–11.

12. Arkhipova, I.R. (2017) Using bioinformatic and phylogenetic approaches to classify transposable elements and understand their complex evolutionary histories. Mob DNA, 8, 19.

13. Huang, X., Zhao, Y., Pu, Q., Liu, G., Peng, Y., Wang, F., Chen, G., Sun, M., Du, F., Dong, J., et al. (2019) Intracellular selection of trans-cleaving hammerhead ribozymes. Nucleic Acids Research, 47, 2514–2522.

14. Radakovic, A., DasGupta, S., Wright, T.H., Aitken, H.R.M. and Szostak, J.W. (2022) Nonenzymatic assembly of active chimeric ribozymes from aminoacylated RNA oligonucleotides. Proceedings of the National Academy of Sciences, 119, e2116840119.

15. Lu, M., Cao, Z., Xiong, L., Deng, H., Ma, K., Liu, N., Qin, Y., Chen, S.-B., Chen, J.-H., Li, Y., et al. (2025) A hammerhead ribozyme selects mechanically stable conformations for catalysis against viral RNA. Commun Biol, 8, 165.

16. Cervera, A., Urbina, D. and de la Peña, M. (2016) Retrozymes are a unique family of non- autonomous retrotransposons with hammerhead ribozymes that propagate in plants through circular RNAs. Genome Biology, 17, 135.

17. Cervera, A. and de la Peña, M. (2020) Small circRNAs with self-cleaving ribozymes are highly expressed in diverse metazoan transcriptomes. Nucleic Acids Research, 48, 5054–5064.

18. Salehi-Ashtiani, K., Lupták, A., Litovchick, A. and Szostak, J.W. (2006) A Genomewide Search for Ribozymes Reveals an HDV-Like Sequence in the Human CPEB3 Gene. Science, 313, 1788–1792.

19. Roth, A., Weinberg, Z., Chen, A.G.Y., Kim, P.B., Ames, T.D. and Breaker, R.R. (2014) A widespread self-cleaving ribozyme class is revealed by bioinformatics. Nat Chem Biol, 10, 56–60.

20. Winkler, W.C., Nahvi, A., Roth, A., Collins, J.A. and Breaker, R.R. (2004) Control of gene expression by a natural metabolite-responsive ribozyme. Nature, 428, 281–286.

21. Ferbeyre, G., Smith, James M. and and Cedergren, R. (1998) Schistosome Satellite DNA Encodes Active Hammerhead Ribozymes. Molecular and Cellular Biology, 18, 3880– 3888.

22. Cloning and characterization of extended hammerheads from a diverse set of caudate amphibians.

23. Przybilski, R., Gräf, S., Lescoute, A., Nellen, W., Westhof, E., Steger, G. and Hammann, C. (2005) Functional Hammerhead Ribozymes Naturally Encoded in the Genome of Arabidopsis thaliana. The Plant Cell, 17, 1877–1885.

24. de la Peña, M. and García-Robles, I. (2010) Ubiquitous presence of the hammerhead ribozyme motif along the tree of life. RNA, 16, 1943–1950.

25. de la Peña, M. (2018) Circular RNAs Biogenesis in Eukaryotes Through Self-Cleaving Hammerhead Ribozymes. In Xiao, J. (ed), Circular RNAs, Advances in Experimental Medicine and Biology. Springer Singapore, Singapore, Vol. 1087, pp. 53–63.

26. De la Peña, M., García-Robles, I. and Cervera, A. (2017) The Hammerhead Ribozyme: A Long History for a Short RNA. Molecules, 22, 78.

27. Hammann, C., Luptak, A., Perreault, J. and Peña, M. de la The ubiquitous hammerhead ribozyme. 10.1261/rna.031401.111.

28. Symons, R.H. (1989) Self-cleavage of RNA in the replication of small pathogens of plants and animals. Trends in Biochemical Sciences, 14, 445–450.

29. Benson, G. (1999) Tandem repeats finder: a program to analyze DNA sequences. Nucleic Acids Res., 27, 573–580.

30. Riccitelli, N.J. and Lupták, A. (2010) Computational discovery of folded RNA domains in genomes and in vitro selected libraries. Methods, 52, 133–140.

31. Lorenz, R., Bernhart, S.H., Höner zu Siederdissen, C., Tafer, H., Flamm, C., Stadler, P.F. and Hofacker, I.L. (2011) ViennaRNA Package 2.0. Algorithms Mol Biol, 6, 26.

32. Rognes, T., Flouri, T., Nichols, B., Quince, C. and Mahé,F. (2016) VSEARCH: a versatile open source tool for metagenomics. PeerJ, 4, e2584.

33. Camacho, C., Coulouris, G., Avagyan, V., Ma, N., Papadopoulos, J., Bealer, K. and Madden, T.L. (2009) BLAST+: architecture and applications. BMC Bioinformatics, 10, 421.

34. Quinlan, A.R. and Hall, I.M. (2010) BEDTools: a flexible suite of utilities for comparing genomic features. Bioinformatics, 26, 841–842.

35. Craig, R.J., Yushenova, I.A., Rodriguez, F. and Arkhipova, I.R. (2021) An Ancient Clade of Penelope-Like Retroelements with Permuted Domains Is Present in the Green Lineage and Protists, and Dominates Many Invertebrate Genomes. Molecular Biology and Evolution, 38, 5005–5020.

36. Jurka, J., Kapitonov, V.V., Pavlicek, A., Klonowski, P., Kohany, O. and Walichiewicz, J. (2005) Repbase Update, a database of eukaryotic repetitive elements. Cytogenet. Genome Res., 110, 462–467.

37. Eddy, S.R. (2011) Accelerated Profile HMM Searches. PLoS Comput Biol, 7, e1002195.

38. Wang, J., Chitsaz, F., Derbyshire, M.K., Gonzales, N.R., Gwadz, M., Lu, S., Marchler, G.H., Song, J.S., Thanki, N., Yamashita, R.A., et al. (2023) The conserved domain database in 2023. Nucleic Acids Res, 51, D384–D388.

39. Rice, P., Longden, I. and Bleasby, A. (2000) EMBOSS: the European Molecular Biology Open Software Suite. Trends Genet, 16, 276–277.

40. Katoh, K. and Standley, D.M. (2013) MAFFT Multiple Sequence Alignment Software Version 7: Improvements in Performance and Usability. Mol Biol Evol, 30, 772–780.

41. Guindon, S., Dufayard, J.-F., Lefort, V., Anisimova, M., Hordijk, W. and Gascuel, O. (2010) New algorithms and methods to estimate maximum-likelihood phylogenies: assessing the performance of PhyML 3.0. Syst Biol, 59, 307–321.

42. Letunic, I. and Bork, P. (2024) Interactive Tree of Life (iTOL) v6: recent updates to the phylogenetic tree display and annotation tool. Nucleic Acids Res, 52, W78–W82.

43. Hellsten, U., Harland, R.M., Gilchrist, M.J., Hendrix, D., Jurka, J., Kapitonov, V., Ovcharenko, I., Putnam, N.H., Shu, S., Taher, L., et al. (2010) The genome of the Western clawed frog Xenopus tropicalis. Science, 328, 633–636.

44. Sela, N., Kim, E. and Ast, G. (2010) The role of transposable elements in the evolution of non-mammalian vertebrates and invertebrates. Genome Biol, 11, R59.

45. Arkhipova, I.R. (2006) Distribution and Phylogeny of Penelope-Like Elements in Eukaryotes. Systematic Biology, 55, 875–885.

46. Chalopin, D., Naville, M., Plard, F., Galiana, D. and Volff, J.-N. (2015) Comparative analysis of transposable elements highlights mobilome diversity and evolution in vertebrates. Genome Biol Evol, 7, 567–580.

47. Hua-Van, A., Le Rouzic, A., Boutin, T.S., Filée, J. and Capy, P. (2011) The struggle for life of the genome’s selfish architects. Biol. Direct, 6, 19.

48. Zhang, H.-H., Peccoud, J., Xu, M.-R.-X., Zhang, X.-G. and Gilbert, C. (2020) Horizontal transfer and evolution of transposable elements in vertebrates. Nat Commun, 11, 1362.

49. Volff, J.-N., Hornung, U. and Schartl, M. (2001) Fish retroposons related to the Penelope element of Drosophila virilis define a new group of retrotransposable elements. Mol Gen Genomics, 265, 711–720.

50. Stephan, W. and Cho, S. (1994) Possible role of natural selection in the formation of tandem-repetitive noncoding DNA. Genetics, 136, 333–341.

51. Scalvenzi, T. and Pollet, N. (2014) Insights on genome size evolution from a miniature inverted repeat transposon driving a satellite DNA. Mol Phylogenet Evol, 81, 1–9.

52. Meštrović, N., Mravinac, B., Pavlek, M., Vojvoda-Zeljko, T., Šatović, E. and Plohl, M. (2015) Structural and functional liaisons between transposable elements and satellite DNAs. Chromosome Res, 23, 583–596.

53. Kramerov, D.A. and Vassetzky, N.S. (2011) Origin and evolution of SINEs in eukaryotic genomes. Heredity, 107, 487–495.

