## Supplemental Figure 1 for "Animal retrozymes are non-autonomous sequences of *Penelope*-like elements"

*Xenopus borealis* REP4 - Rtz_CM044436.1.4 – (544/273)

#### ZL1-10 min

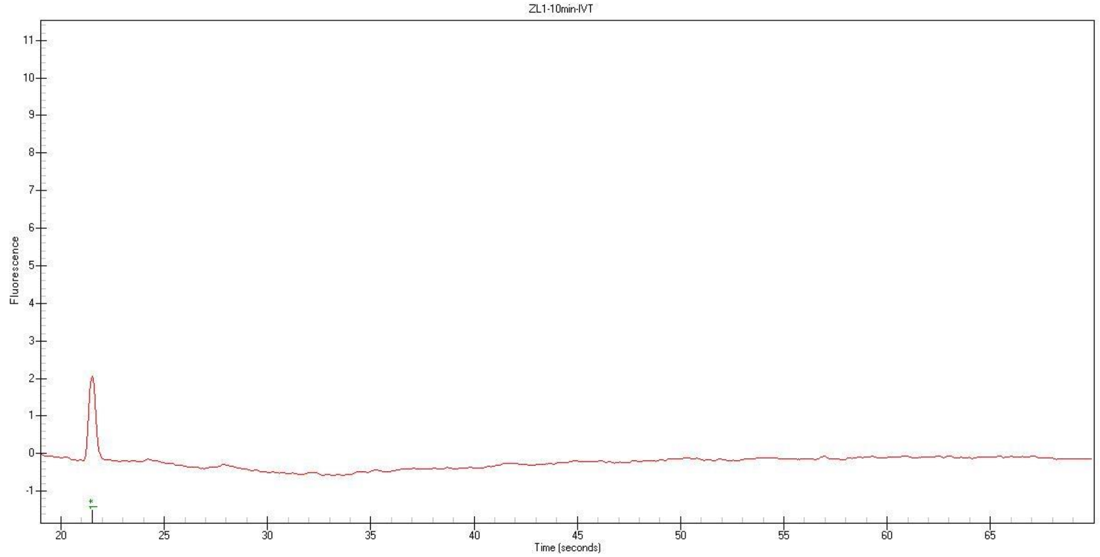


#### ZL1-30 min

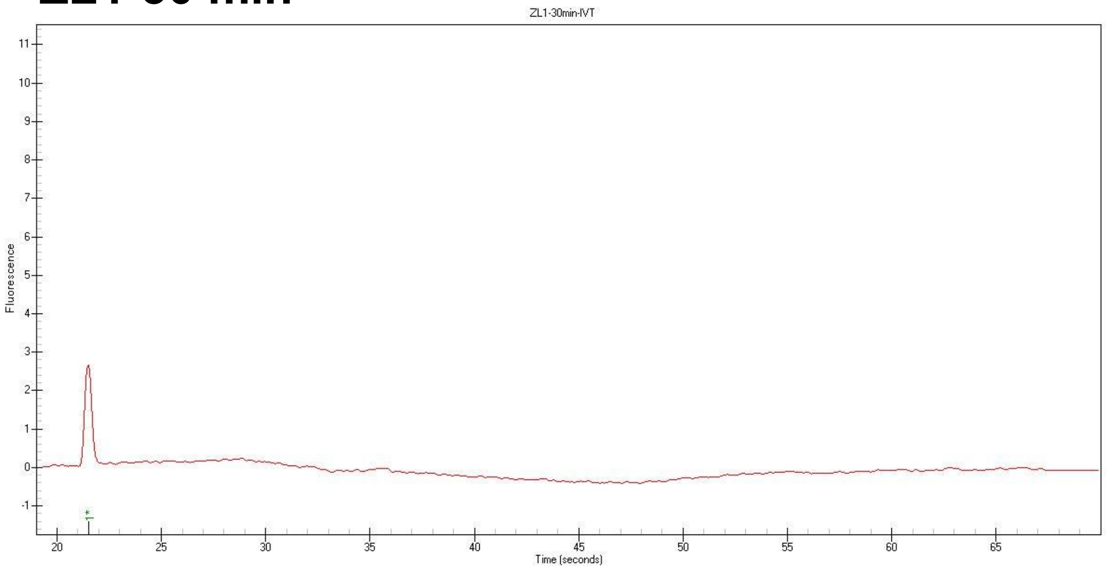


#### ZL1-60 min

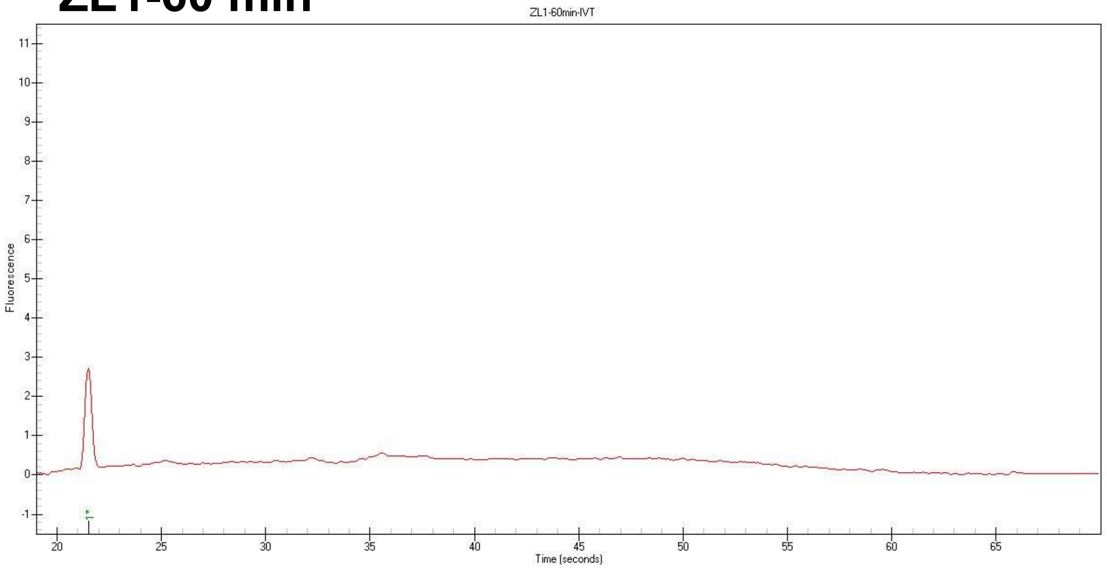


#### ZL1-100 min

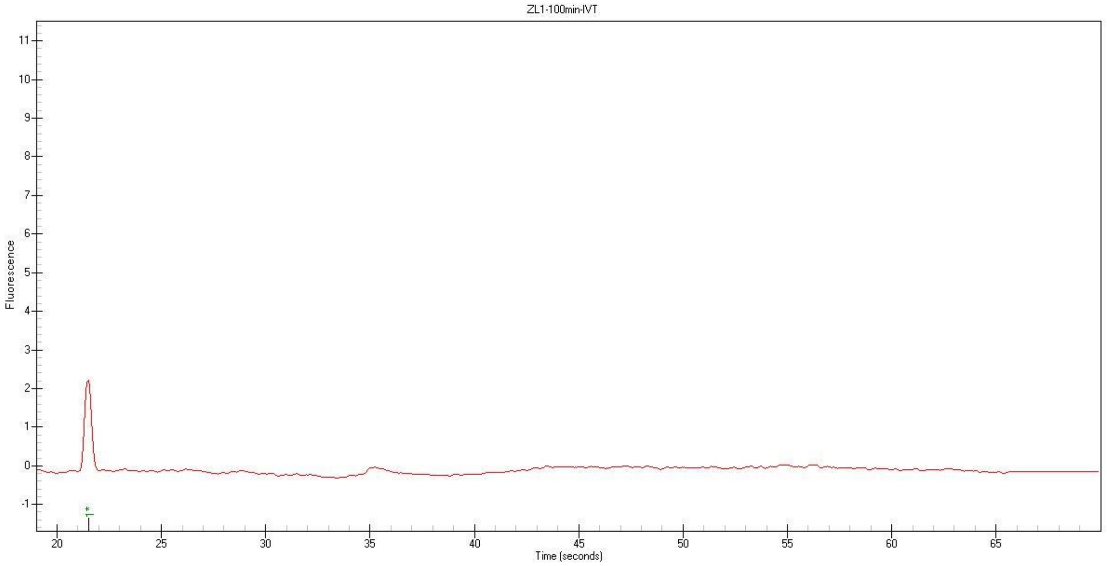

*Xenopus borealis* Penelope-6_XT - Rtz_CM044442.1.5– (221/-)

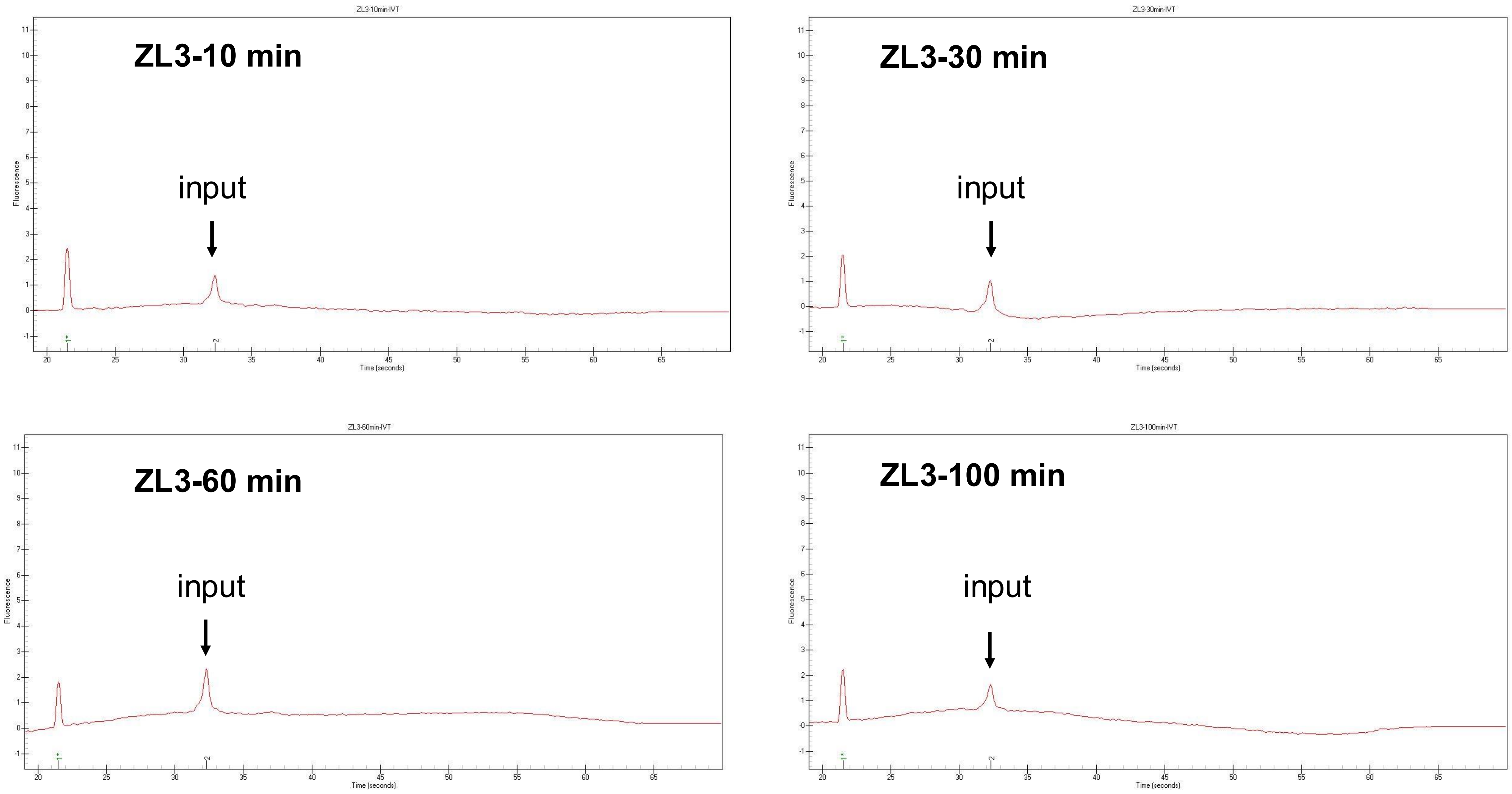

*Xenopus tropicalis* REP4 - Rtz_NC_030678.2.4– (666/-)

**ZL5-10 min**

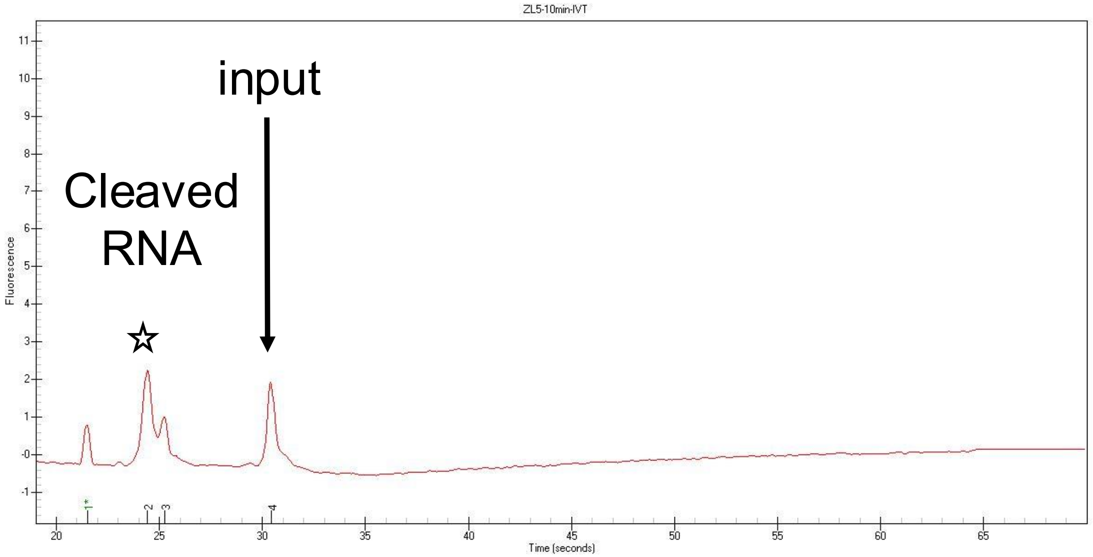


**ZL5-30 min**

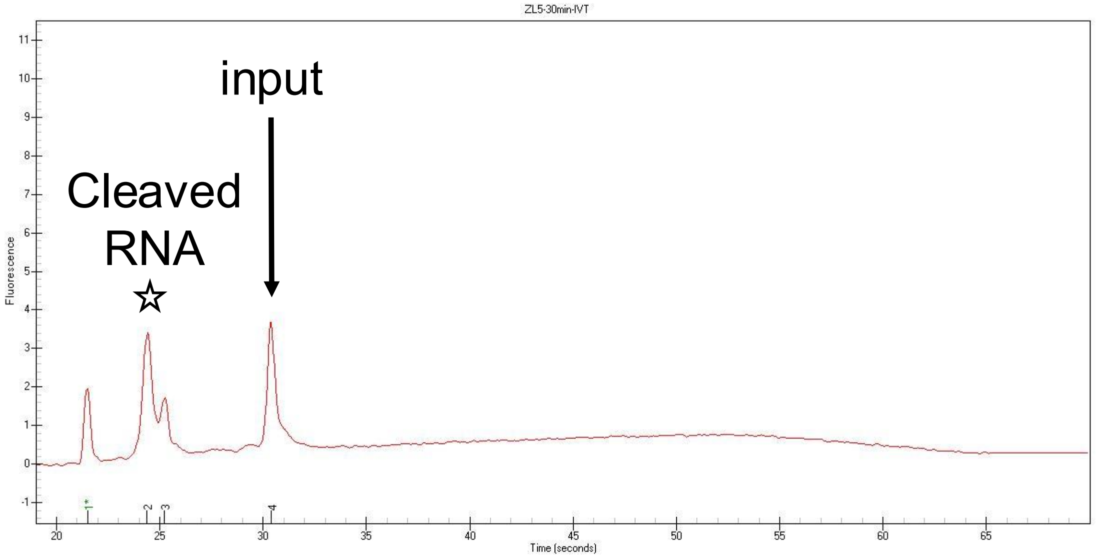


**ZL5-60 min**

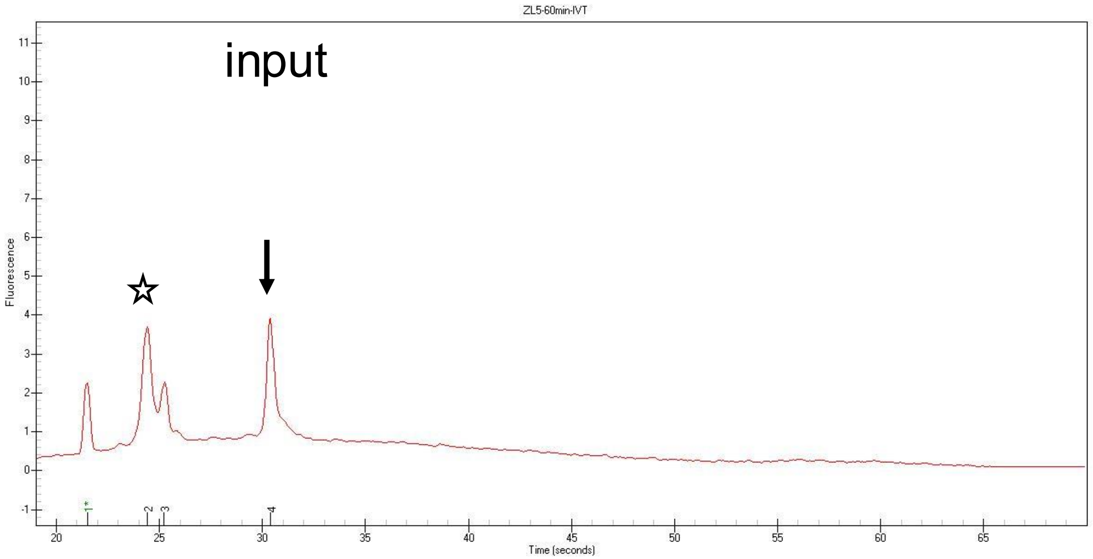


**ZL5-100 min**

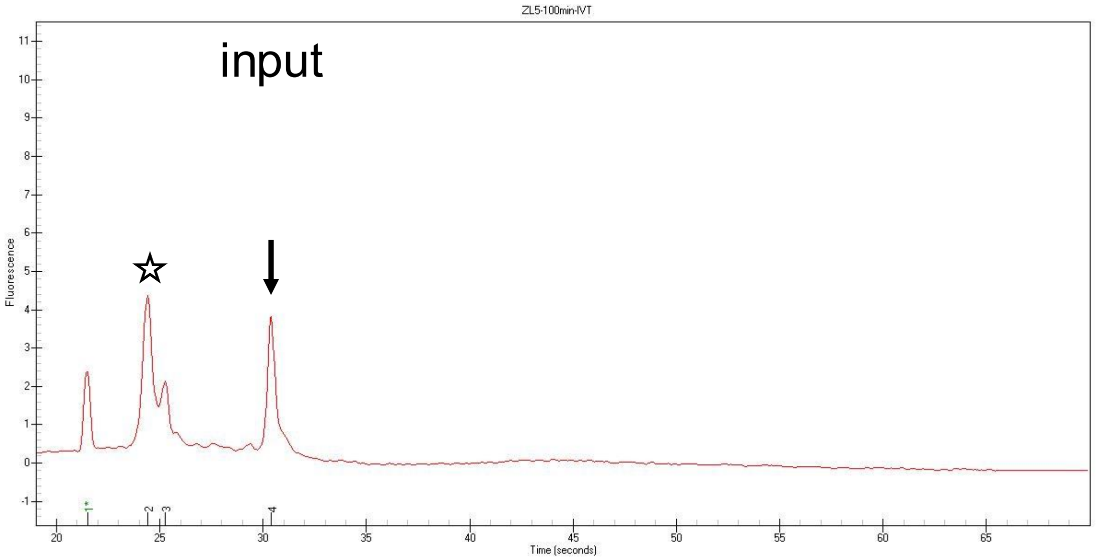

*Xenopus tropicalis* Penelope-6_XT - Rtz_NC_030682.2.3 - (291/-)

#### ZL7-10 min

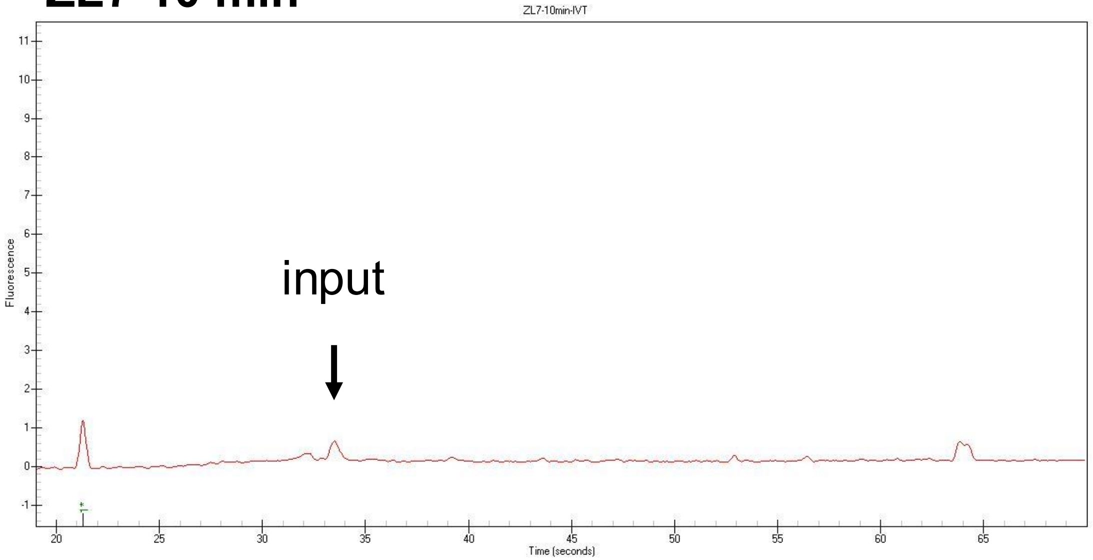


#### ZL7-30 min

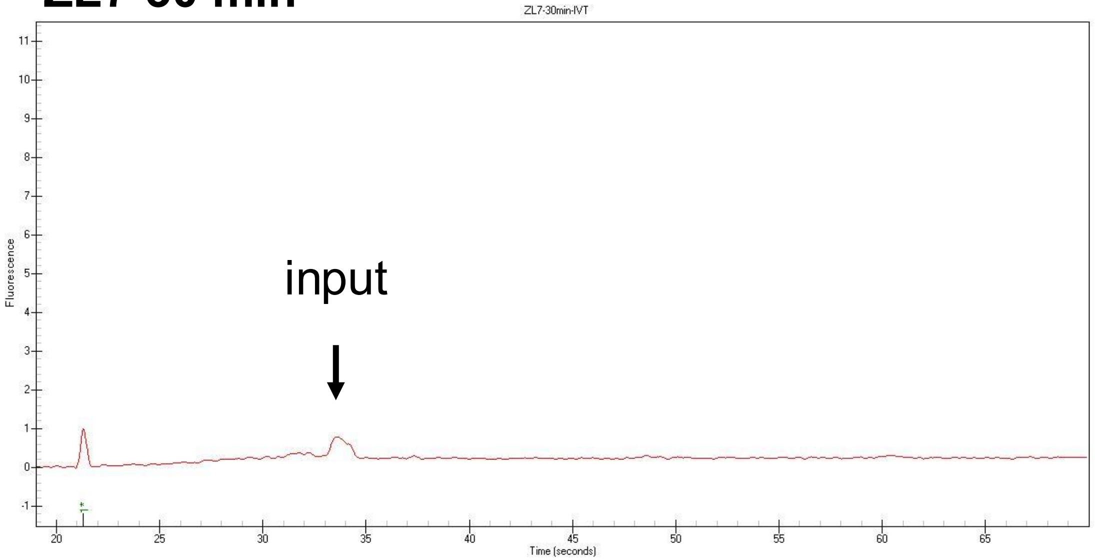


#### ZL7-60 min

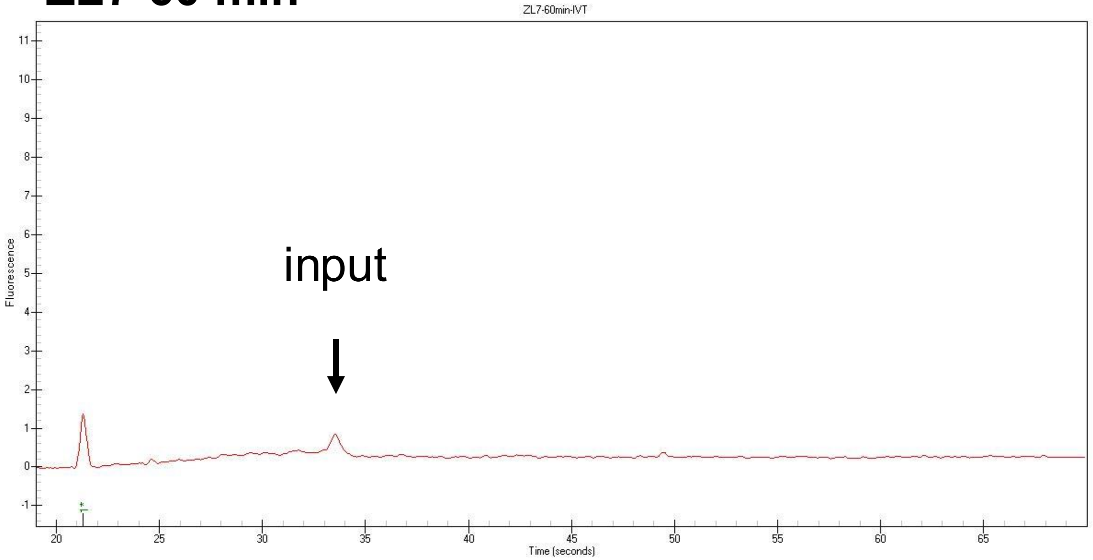


#### ZL7-100 min

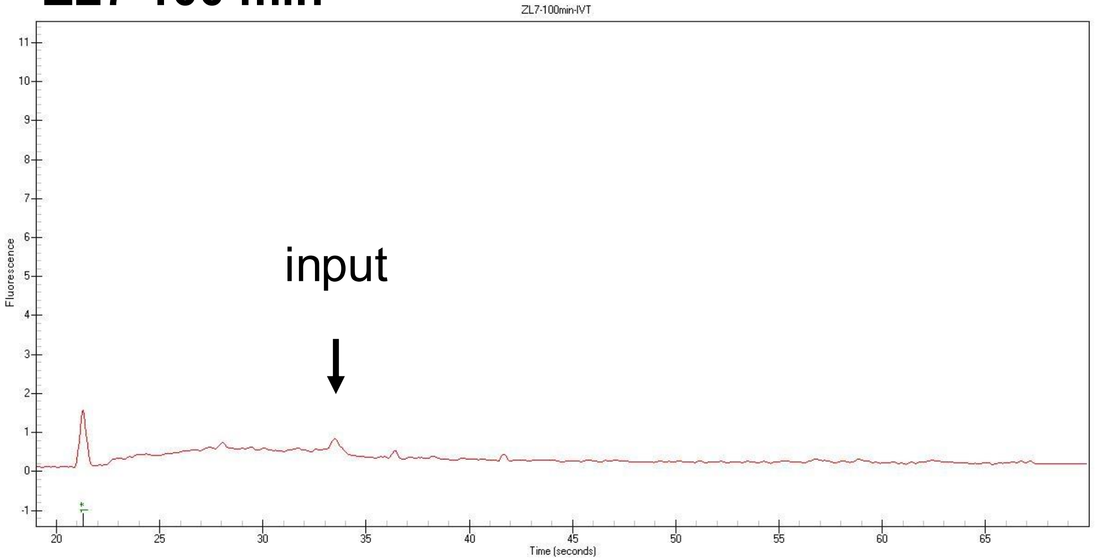

*Xenopus tropicalis* REP4 - Rtz_NC_030685.2.6- (472/275

**ZL9-10 min**

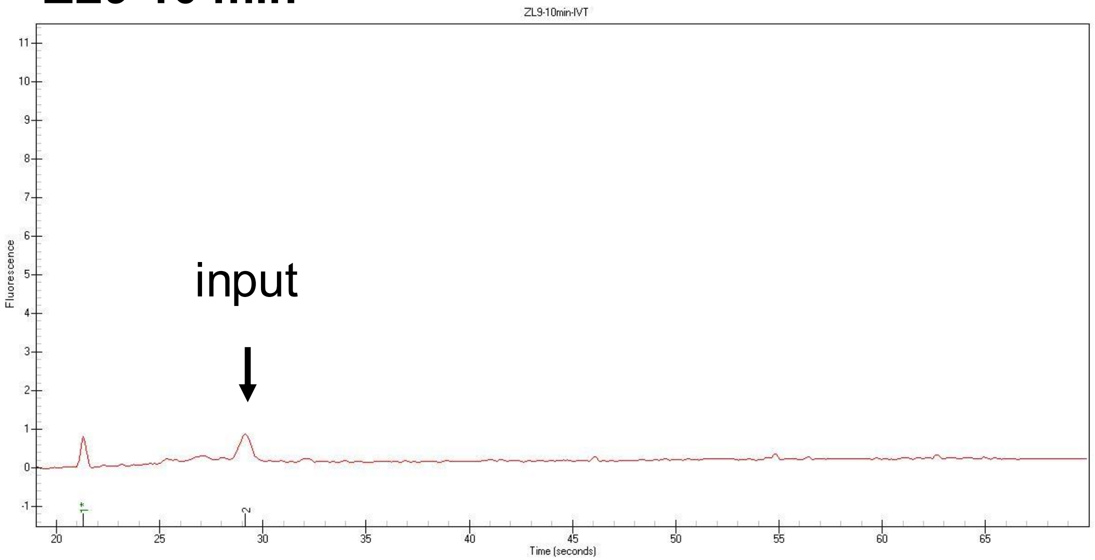


**ZL9-30 min**

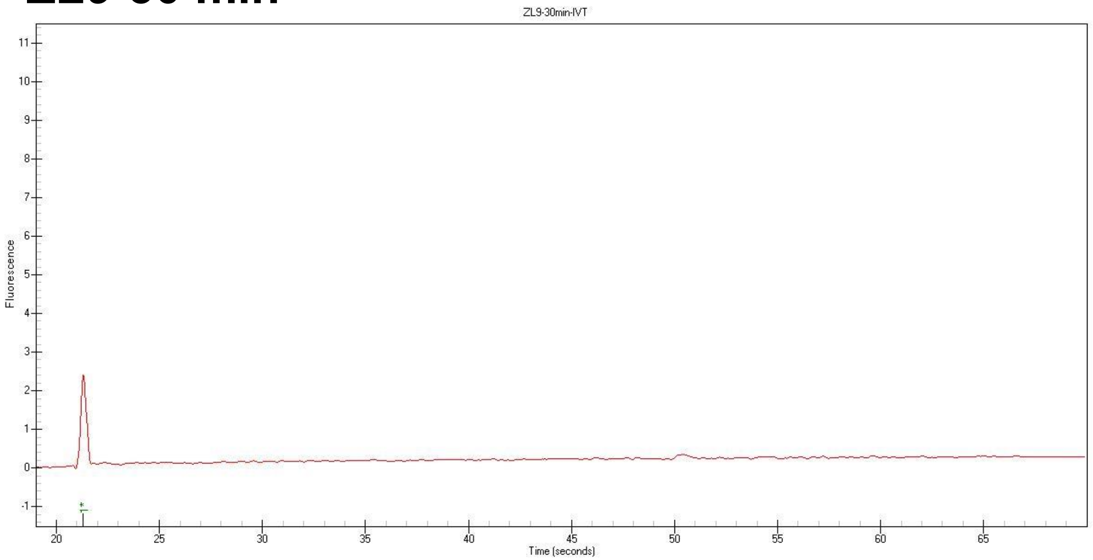


**ZL9-60 min**

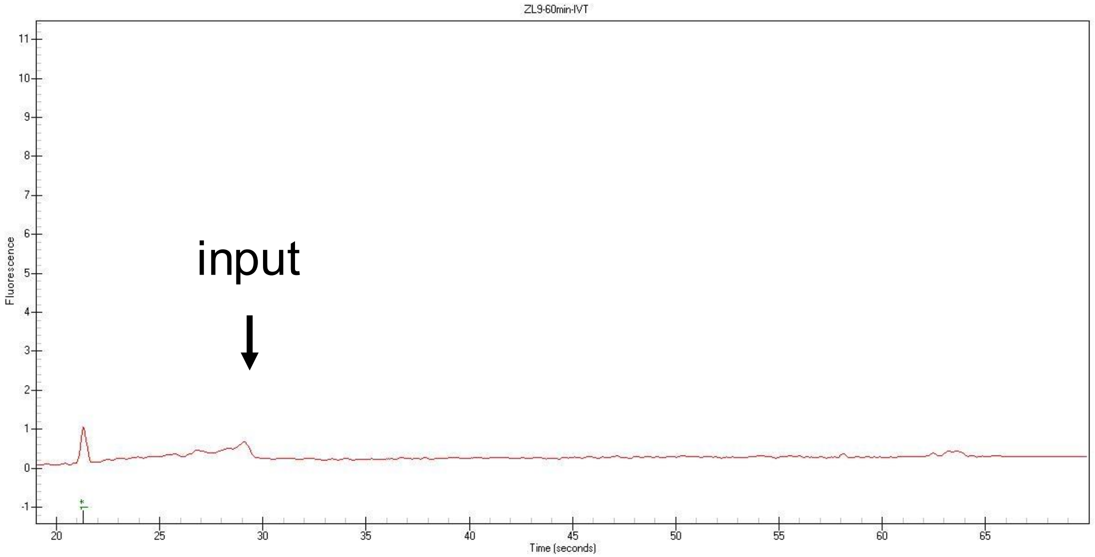


**ZL9-100 min**

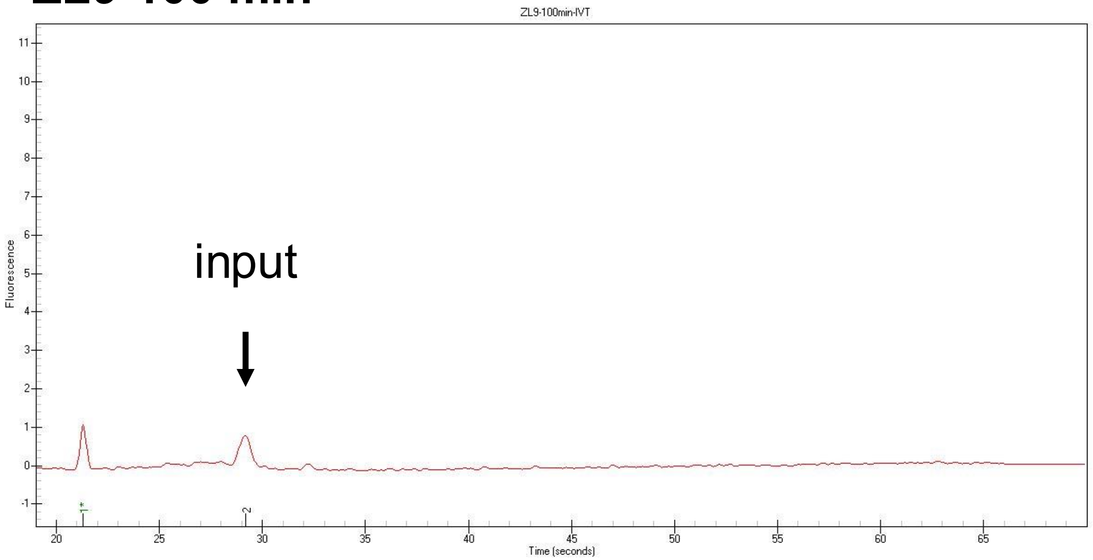

*Xenopus tropicalis Neptune-46_XT Rtz_NC_030685.2.8*

**ZL11-10 min**

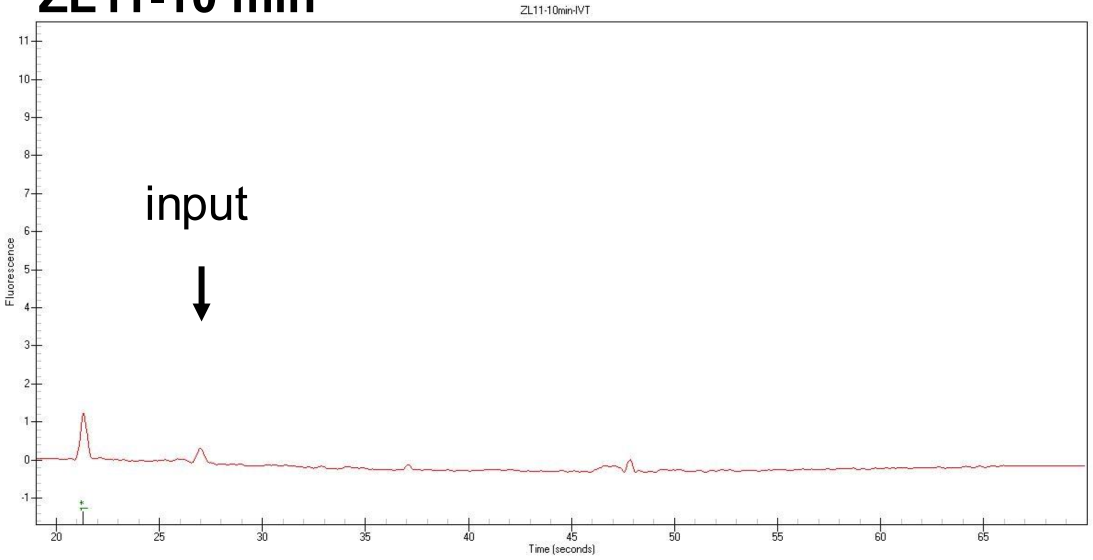


**ZL11-30 min**

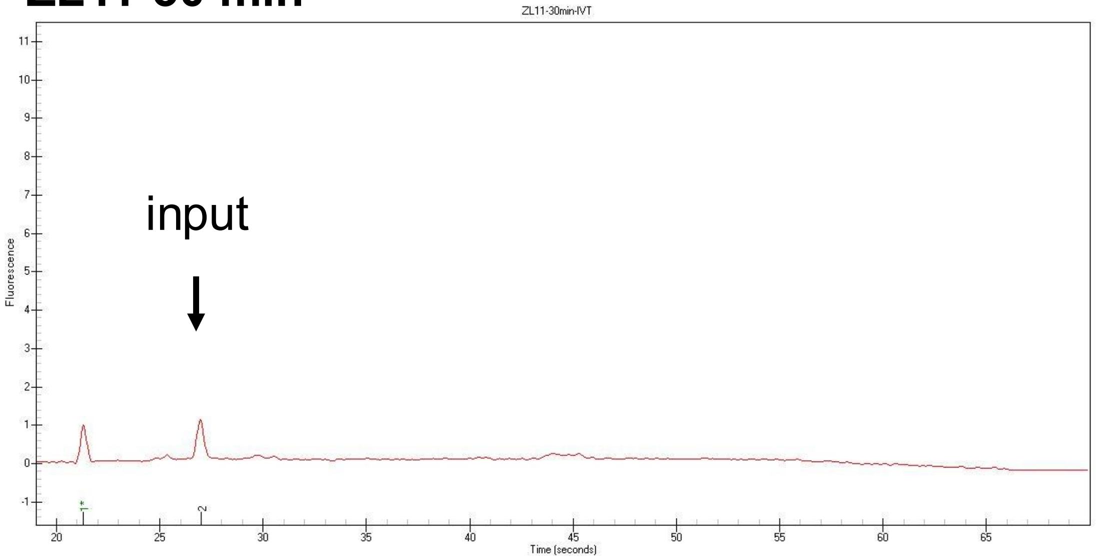


**ZL11-60 min**

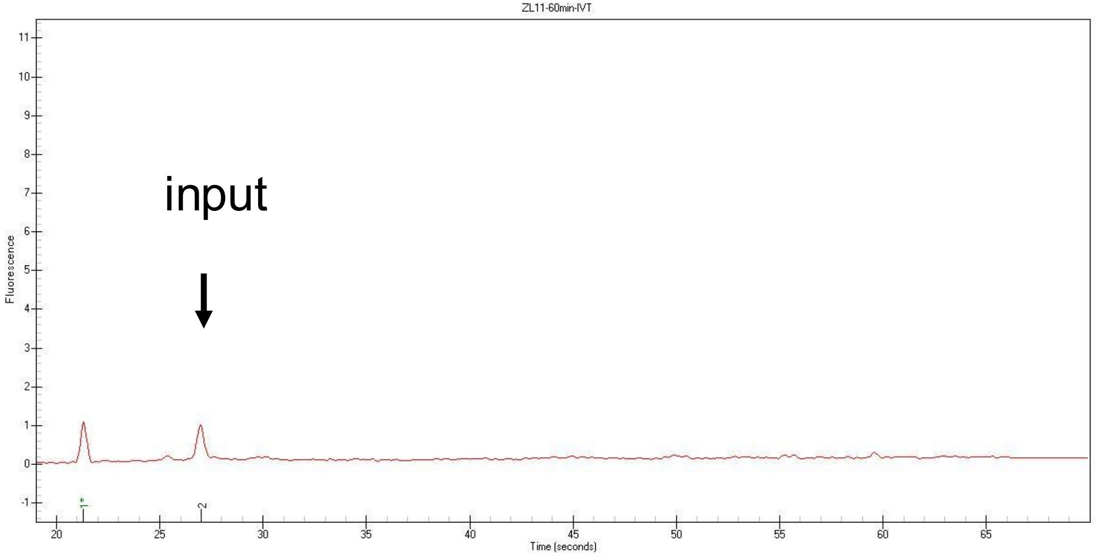


**ZL11-100 min**

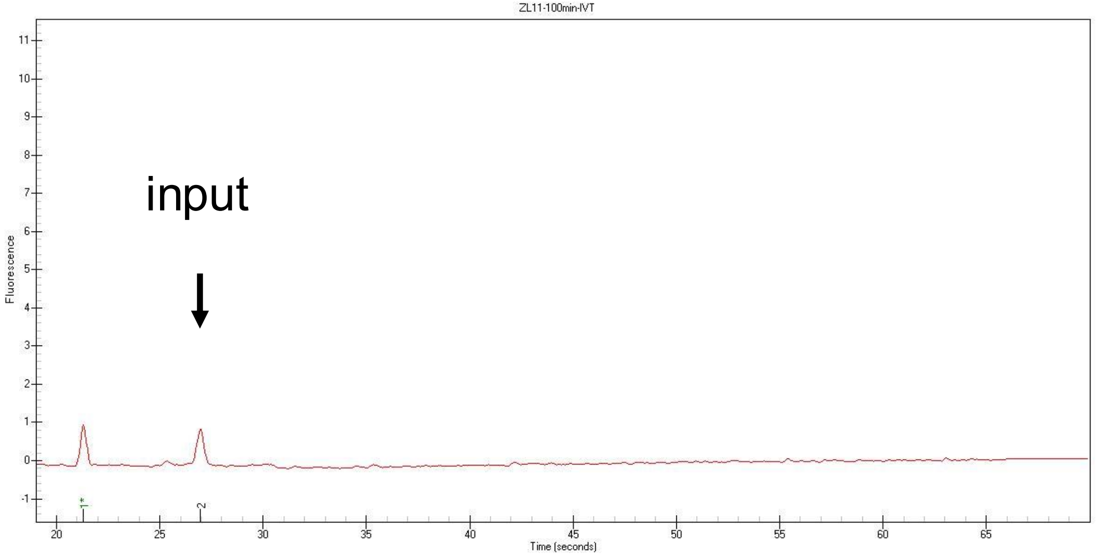

*Xenopus tropicalis Penelope-6_XT Rtz_NC_030686.2.2*

**ZL13-10 min**

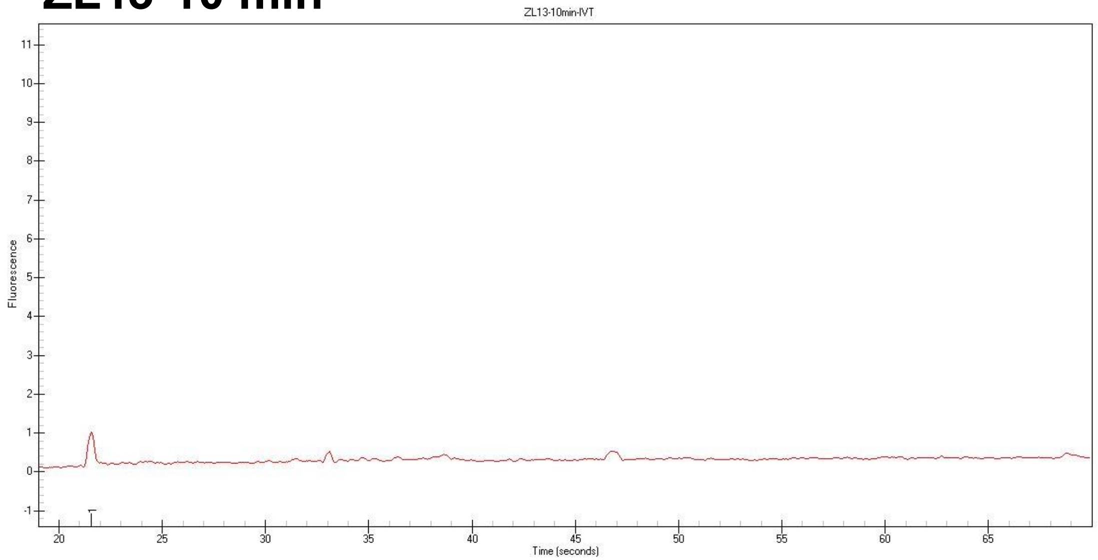


**ZL13-30 min**

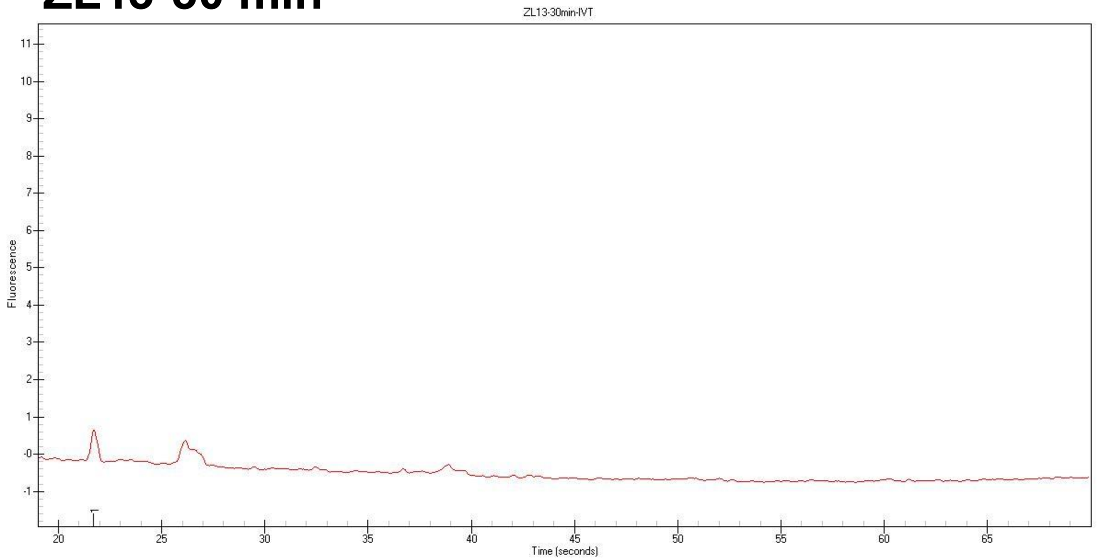


**ZL13-60 min**

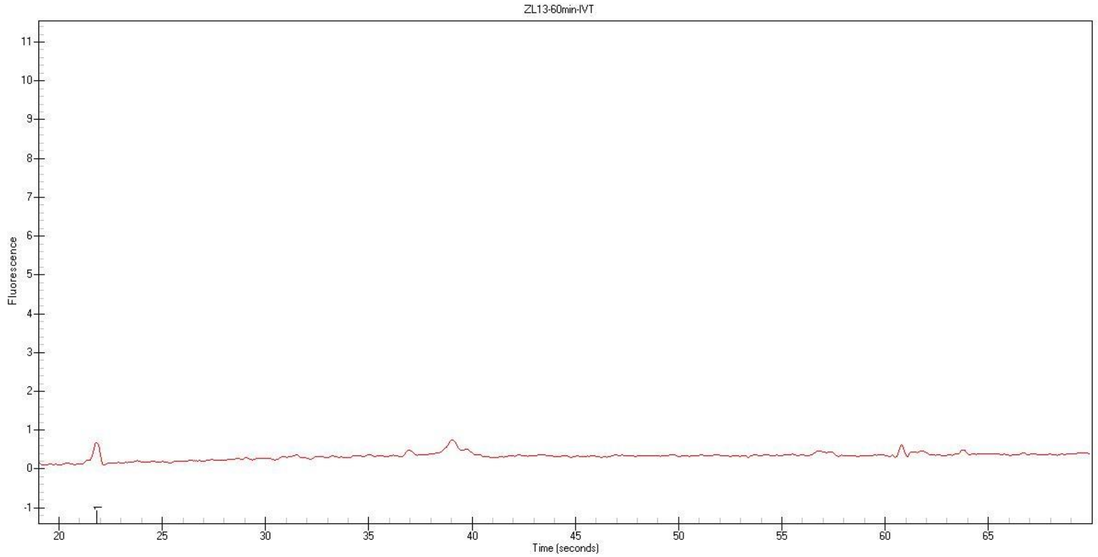


**ZL13-100 min**

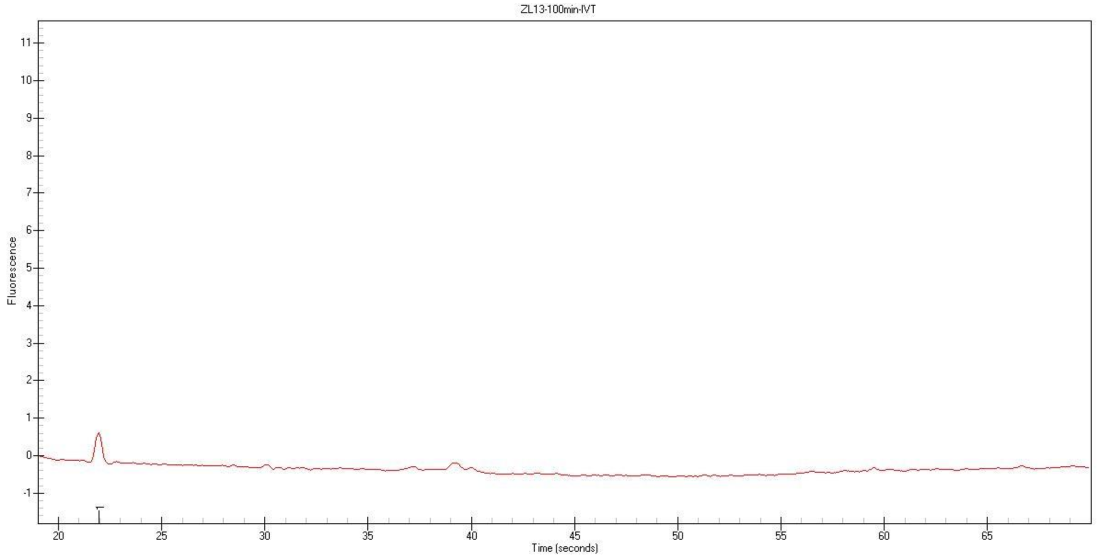

### *Xenopus laevis* Neptune-9_XL Rtz_NC_054382.1.10

#### ZL17-10 min

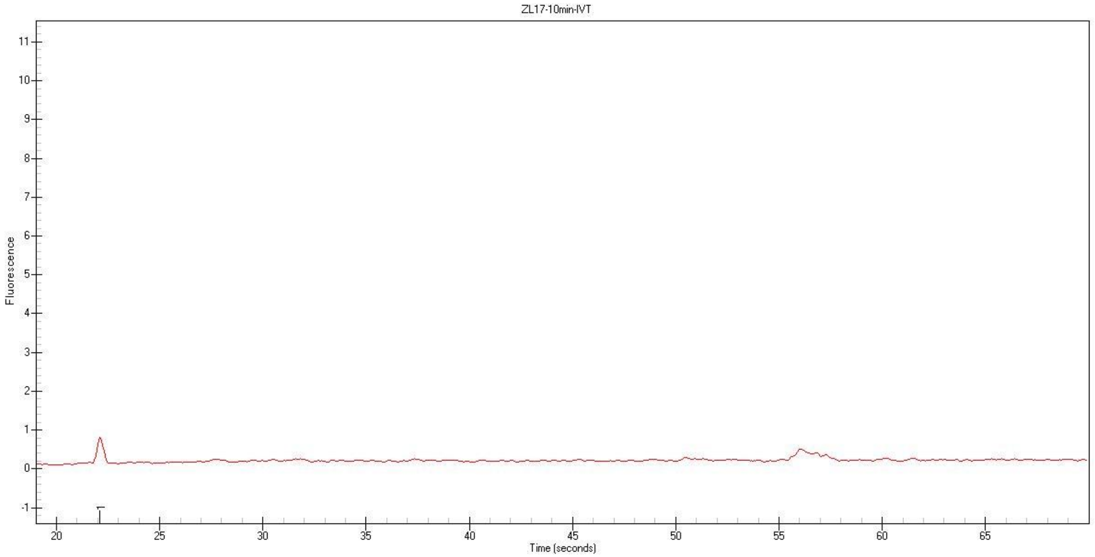


#### ZL17-30 min

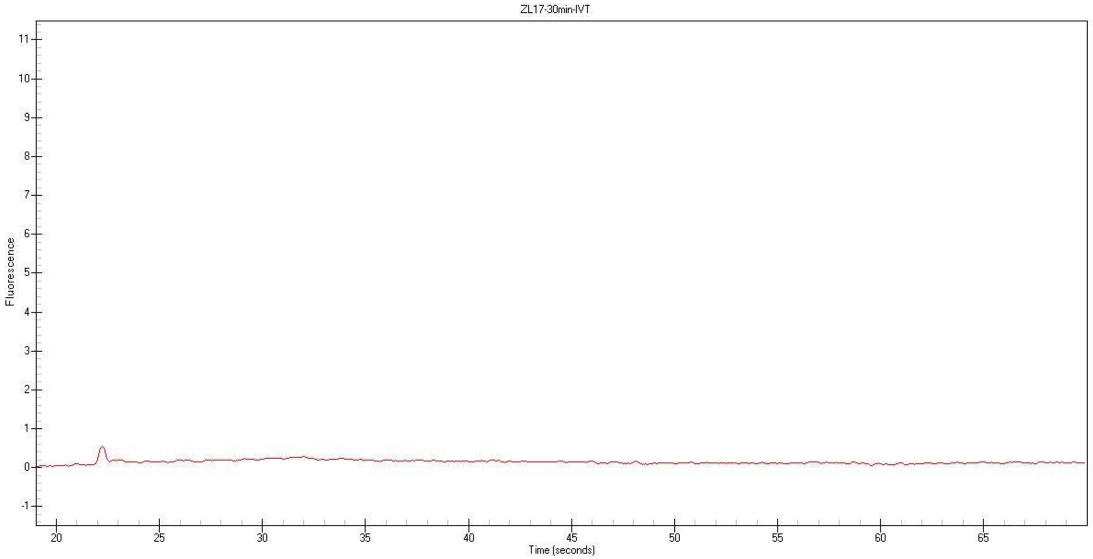


#### ZL17-60 min

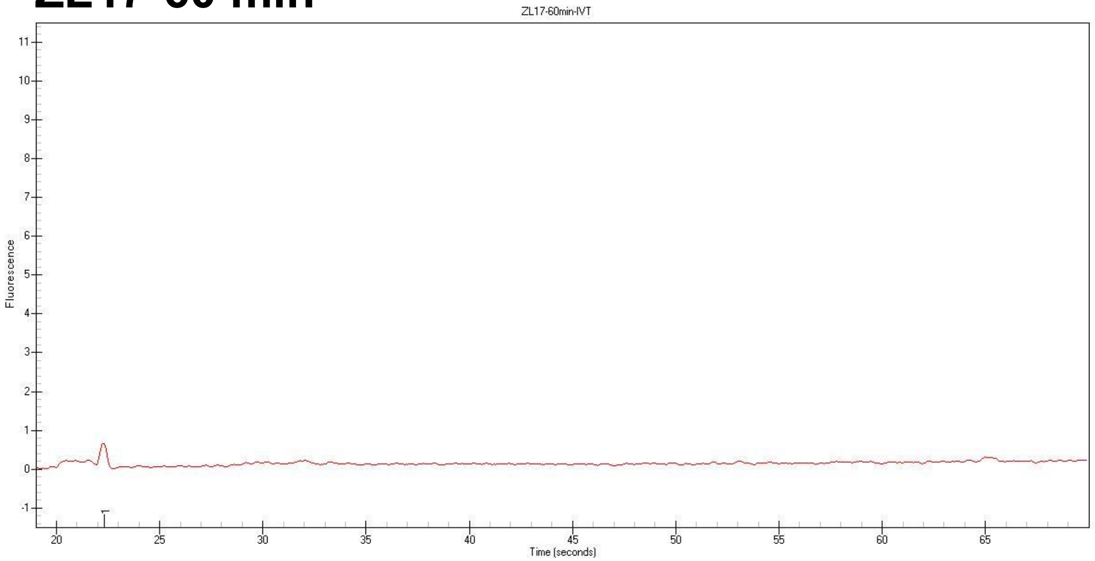


#### ZL17-100 min

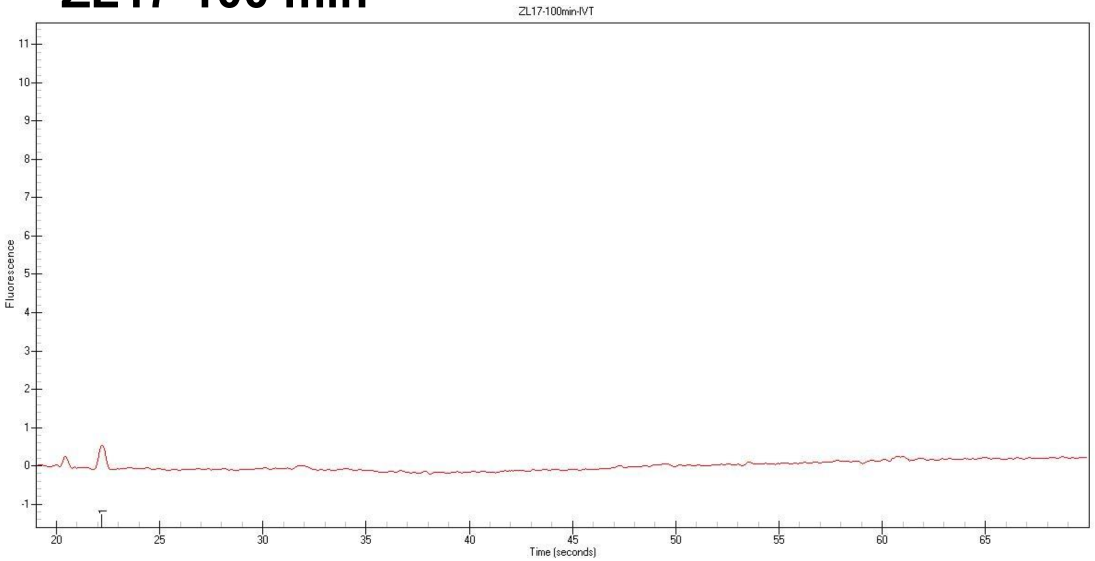

*Xenopus laevis* Neptune Rtz_NC_054387.1.4

**ZL19-10 min**

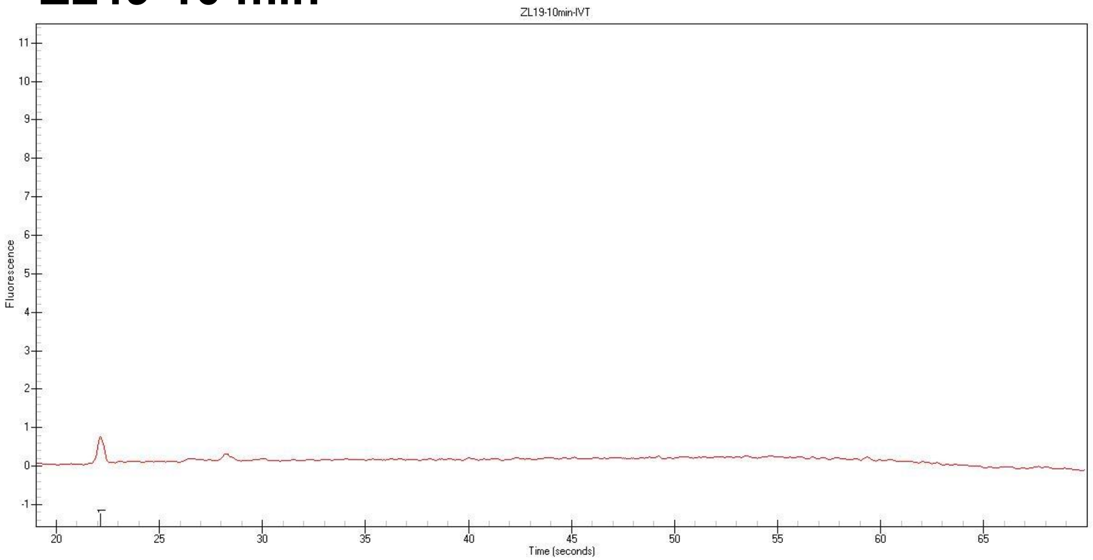


**ZL19-30 min**

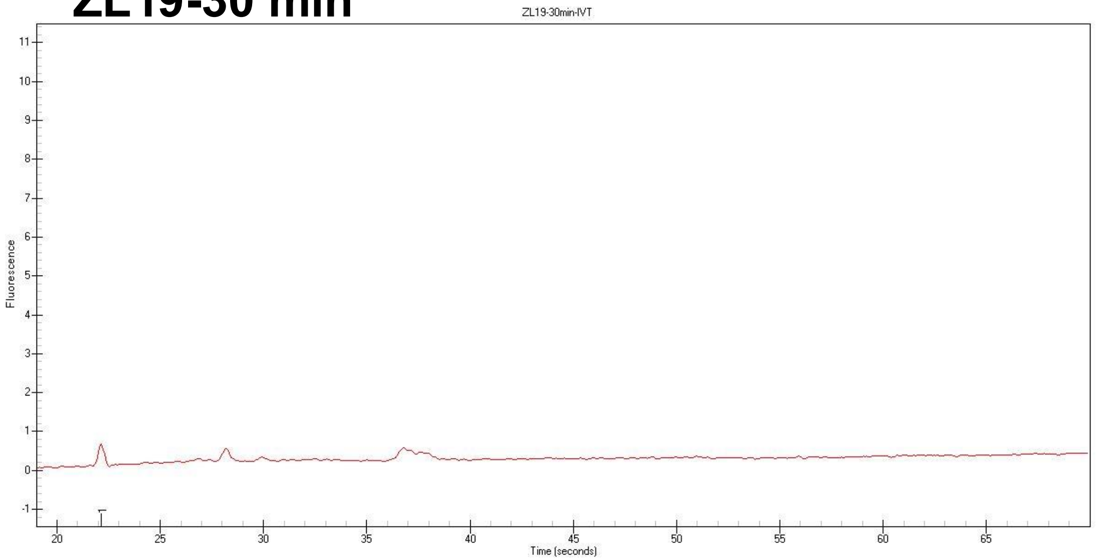


**ZL19-60 min**

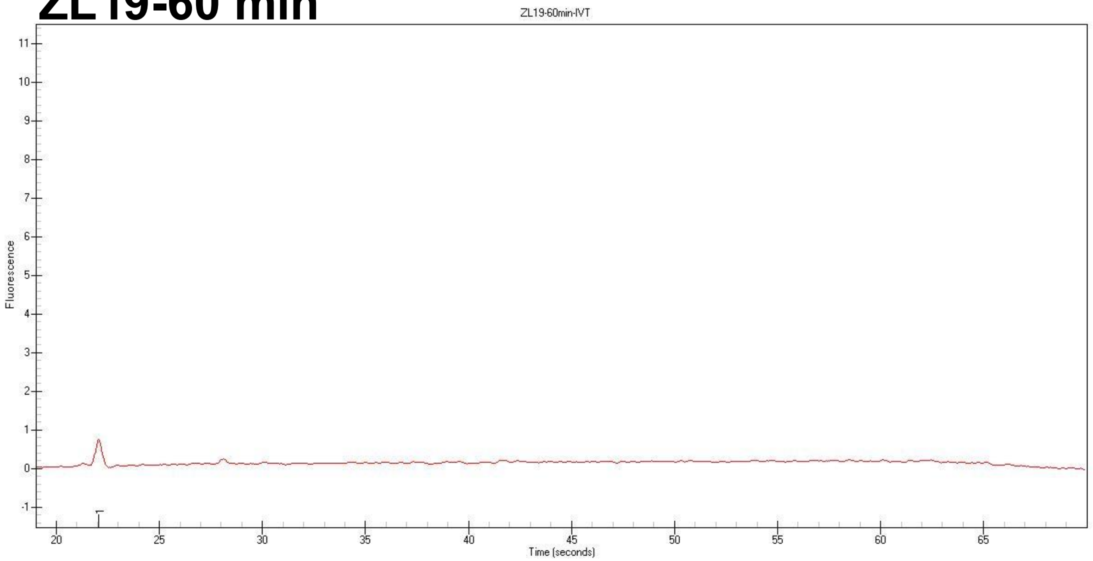


**ZL19-100 min**

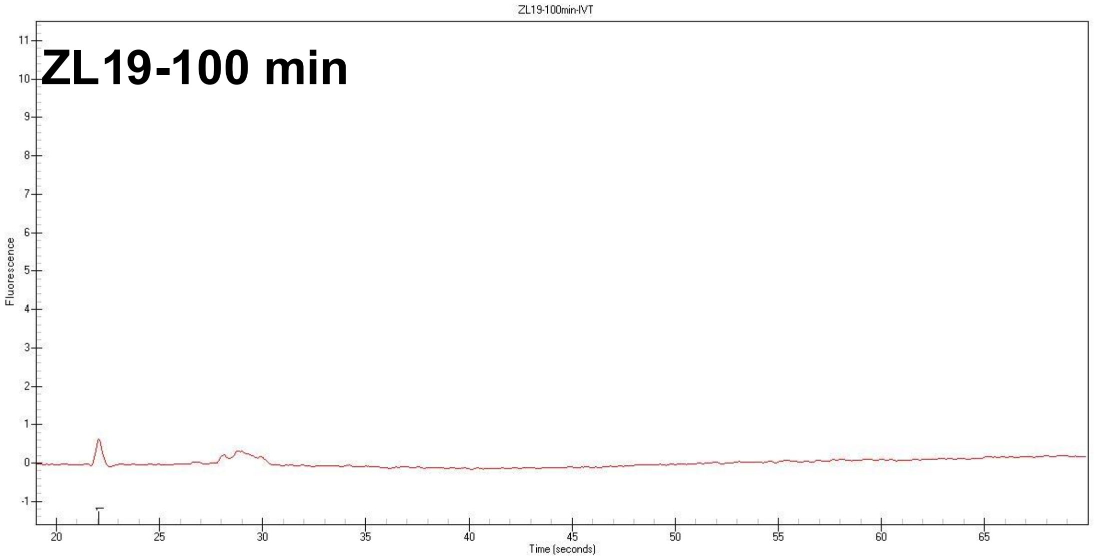
